# A robotic platform for fluidically-linked human body-on-chips experimentation

**DOI:** 10.1101/569541

**Authors:** Richard Novak, Miles Ingram, Susan Clauson, Debarun Das, Aaron Delahanty, Anna Herland, Ben M. Maoz, Sauveur S. F. Jeanty, Mahadevabharath R. Somayaji, Morgan Burt, Elizabeth Calamari, Angeliki Chalkiadaki, Alexander Cho, Youngjae Choe, David Benson Chou, Michael Cronce, Stephanie Dauth, Toni Divic, Jose Fernandez-Alcon, Thomas Ferrante, John Ferrier, Edward A. FitzGerald, Rachel Fleming, Sasan Jalili-Firoozinezhad, Thomas Grevesse, Josue A. Goss, Tiama Hamkins-Indik, Olivier Henry, Chris Hinojosa, Tessa Huffstater, Kyung-Jin Jang, Ville Kujala, Lian Leng, Robert Mannix, Yuka Milton, Janna Nawroth, Bret A. Nestor, Carlos F. Ng, Blakely O’Connor, Tae-Eun Park, Henry Sanchez, Josiah Sliz, Alexandra Sontheimer-Phelps, Ben Swenor, Guy Thompson, George J. Touloumes, Zachary Tranchemontagne, Norman Wen, Moran Yadid, Anthony Bahinski, Geraldine A. Hamilton, Daniel Levner, Oren Levy, Andrzej Przekwas, Rachelle Prantil-Baun, Kevin K. Parker, Donald E. Ingber

## Abstract

Here we describe of an ‘Interrogator’ instrument that uses liquid-handling robotics, a custom software package, and an integrated mobile microscope to enable automated culture, perfusion, medium addition, fluidic linking, sample collection, and *in situ* microscopic imaging of up to 10 Organ Chips inside a standard tissue culture incubator. The automated Interrogator platform maintained the viability and organ-specific functions of 8 different vascularized, 2-channel, Organ Chips (intestine, liver, kidney, heart, lung, skin, blood-brain barrier (BBB), and brain) for 3 weeks in culture when fluidically coupled through their endothelium-lined vascular channels using a common blood substitute medium. When an inulin tracer was perfused through the multi-organ Human Body-on-Chips (HuBoC) fluidic network, quantitative distributions of this tracer could be accurately predicted using a physiologically-based multi-compartmental reduced order (MCRO) *in silico* model of the experimental system derived from first principles. This automated culture platform enables non-invasive imaging of cells within human Organ Chips and repeated sampling of both the vascular and interstitial compartments without compromising fluidic coupling, which should facilitate future HuBoc studies and pharmacokinetics (PK) analysis *in vitr*o.

---

Vascularized human Organ Chips are microfluidic cell culture devices containing separate vascular and parenchymal compartments lined by living human organ-specific cells that recapitulate the multicellular architecture, tissue-tissue interfaces, and relevant physical microenvironments of key functional units of living organs, while providing vascular perfusion *in vitro*^1,2^. The growing recognition that animal models do not effectively predict drug responses in humans^3–5^ and the related increase in demand for *in vitro* human toxicity and efficacy testing, has led to pursuit of time-course analyses of human Organ Chip models and fluidically linked, multi-organ, HuBoC systems that recapitulate organ-level functions to facilitate studies of drug pharmacokinetics and pharmacodynamics (PK/PD) *in vitro*^6–11^.

An automated experimental system that can meet these goals needs to be highly multi-functional and must ideally enable fluid handling and sample collection, perfusion of fluid through multiple linked microfluidic Organ Chip devices, and tissue imaging, all within a controlled temperature, humidity, and CO_2_ environment. Custom assemblies using syringe pumps^12^, peristaltic pumps^13^, micropumps^11,14,15^, and gravity feed^10,16,17^, including several commercial platforms^18,19^, can be used to perfuse and link microfluidic tissue and organ culture inside incubators; however, these systems do not facilitate the features necessary for complex HuBoC experimentation. In particular, these systems offer limited ability to sample the different biological compartments or the capacity to reconfigure the system so that the same platform can be used for multiple experimental designs. Other HuBoC systems have used manual or automated transfer of fluid flow between multiple microfluidic culture systems using gravity feed^10,16^ or they relied on perfusion through integrated microfluidic networks controlled by microvalve pumps^14,19,20^. But in these systems the shared medium containing drugs was transferred directly from one parenchymal tissue type to another without passing across a vascular endothelium as normally occurs in vivo. This lack of endothelial barriers is a significant limitation in PK studies where the endothelium can contribute significantly to drug absorption, distribution, metabolism, excretion (ADME) and toxicity^21,22^.

Here, we present a new approach to linking different Organ Chips for HuBoC studies that employs liquid-handling robotics to overcome these limitations. Specifically, this approach relies on the automatic and regular robotic transfer of liquid samples between individual Organ Chip that are each perfused continuously. The liquid transfers act to replace direct fluidic plumbing (e.g. through tubing or microfluidic channels), thereby avoiding various engineering complexities, such as priming, washing, and a propensity to collect bubbles, while providing full reconfigurability and software-based control. Perfusion pumps are assigned to each of the Organ Chips to ensure that they experience continuous fluid flow independently of the action of the robotic fluid transfer system.

To include liquid-handling robotics and individual Organ-Chip perfusion in a format that fits within standard tissue-culture incubators, we developed a custom, modular platform for culture and analysis, which we call an ‘Interrogator’ instrument. In addition to the automated culture, perfusion and fluidic coupling, we incorporated *in situ* imaging using a software-controlled mobile microscope. We also developed a, Interrogator control environment that permits the on-demand reconfiguration of experimental protocols, routing of drug administration, media replenishment, and analyte sampling, as well as Organ Chip addition, removal, and exchange. The system supports vascularized Organ-Chips composed of poly-dimethlysiloxane (PDMS) or polycarbonate (PC) and that contain two parallel, continuously perfused microchannels separated by a porous membrane, which are seeded with living human organ-specific parenchymal cells and vascular endothelial cells on either side of the membrane to create a tissue-tissue interface^23,24^. The dimensions of the channels, perfusion rate, medium types, and protein coatings can be varied to create organ-specific designs that optimize surface area, or in the case of skin, that lacks an upper surface of the top channel to permit experimental access to the surface of the epidermis (**Supplementary Fig. S1**). The endothelium-lined vascular channel in each Organ Chip is perfused with a common ‘blood substitute’ universal medium (**See method section**), while each parenchymal channel is perfused with organ-specific medium or exposed to an air-liquid interface (in the case of lung and skin) to more accurately emulate organ-level physiology and pathophysiology^1,25–33^. The Organ Chips also contain hollow side chambers that run in parallel to the central culture channels through which cyclic suction can be applied to mechanically stretch and relax the cultured tissue-tissue interface, and thereby mimic physiological tissue motions that occur *in vivo* (e.g., breathing in lung, peristalsis in gut)^28,29,34,35^. Our studies demonstrate that the automated Interrogator instrument can be used to maintain multiple human vascularized Organ Chips for weeks in culture when fluidically linked. Using the Interrogator, it is also possible to emulate physiological systemic transport of small molecules between organs through a common endothelium-lined vasculature and across the endothelium-parenchymal tissue interface of each organ, which is a major contributor to drug PK behavior^2,22^.

## RESULTS

### Automated Interrogator instrument design and operation

The automated Interrogator platform was designed to be small (45 × 45 × 45 cm) so that it can fit within a standard tissue culture incubator and provide continuous perfusion of up to 10 Organ Chips for HuBoC studies using an integrated custom peristaltic pump module and fluidic coupling via their vascular channel reservoirs (Fig. 1). The Interrogator also enables automated sampling and dosing of chips by transferring medium samples to and from wells of standard well plates housed in the instrument and the inlets and outlets of the Organ Chips using a robot fluid handling device. The instrument, which was fabricated in-house, permits automated transfers of controlled fluid volumes (>50 μL) between arbitrary chips and reservoirs in a programmable manner for organ-organ linking as well as for experimental sampling (**Supplementary Movies 1 and 2**). The system can emulate different drug delivery methods, including intravenous, oral, inhalational, or transdermal, by delivering compounds either to an arteriovenous (AV) reservoir that links to the vascular channels of all chips or to the parenchymal channels of Gut, Lung or Skin Chips, respectively. A server box lies outside of the incubator to provide networked connection from a webapp interface. Low-level electronics, including motorized axis controls and camera, are located within the incubator and can withstand the high humidity and CO_2_ atmosphere. A standard USB cable and power cable are the only connections necessary for operation.

**Figure 1.**
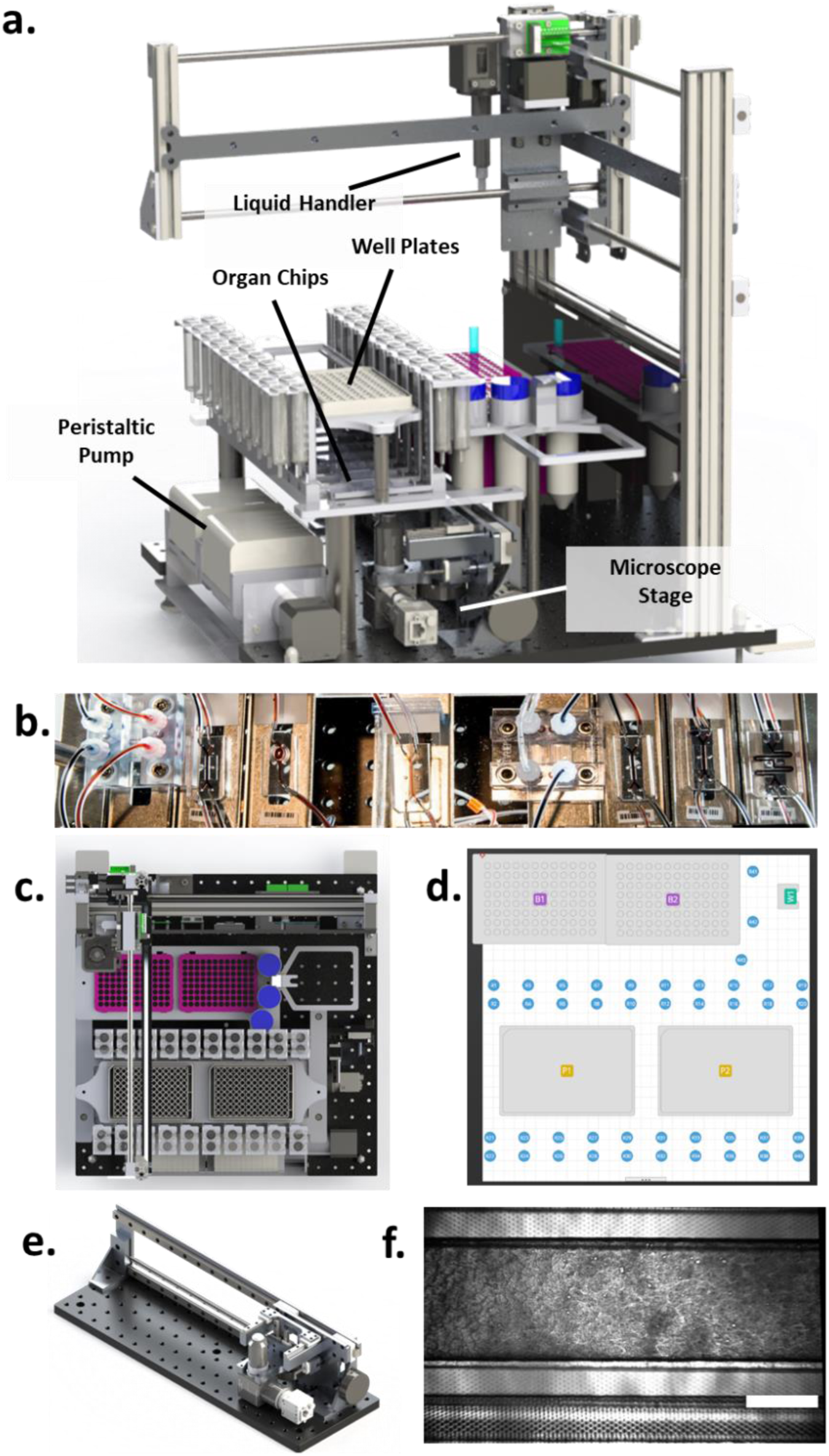
**(a)** Rendering of the Interrogator CAD model. The system is comprised of a 3-axis motion system, automated liquid handler, peristaltic pump, and custom microscope stage that allows for the continuous perfusion, linking and image analysis of organ-on-chip models. **(b)** Organ Chips are placed between inlet and outlet reservoirs in modular docks below the main fluid handling deck. Top view of the instrument deck layout in CAD **(c)** and within the graphical user interface **(d)**. Component positions are determined from CAD then used to build a virtual deck in the control software. **(e)** Rendering of the microscope module, showing a compact optical path microscope mounted on a 3-axis positioning stage that is integrated into the Interrogator for real-time imaging and inspection of Organ Chips. **(f)** Micrograph of a Gut Chip channel showing villus-like structures.

The Interrogator instrument is comprised of multiple subsystems, including a 3-axis motion system, automatic liquid handler, computer-controlled peristaltic pump, and a microscope imaging module (Fig. 1a). The configuration described here is designed to support up to 10 different vascularized (2-channel) Organ Chips (Fig. 1b), while the deck layout is customizable and reconfigurable (Fig. 1c,d). Each Organ Chip module features a completely independent peristaltic perfusion system, consisting of two inlet reservoirs, all tubing connections, and two outlet reservoirs. Automated linking between Organ Chips can be achieved by either direct and repeated pipetting of discrete volumes of medium from one chip outlet to the inlet of another chip, or through an intermediate reservoir. This reconfigurability and modularity enables the user to change chip linkage order and experimental protocols without the need to physically move Organ Chips or change tubing connections, which can restrict use of existing HuBoC systems. The Instrument also contains a standalone microscope module (Fig. 1e,f) consisting of a three-axis miniaturized stage and a compact microscope with camera, which enables on-demand imaging of the Organ Chips without requiring cessation of fluidic coupling or removal from the incubator.

The custom software environment developed to control the Interrogator provides a graphical user interface for instrument deck setup, configuration of reservoirs and other consumables, experimental design and execution, and real-time experimental error-checking to preempt costly programming errors (**Supplementary Fig. S2**). Complex operations, such as multi-well sampling and serial dilutions, are handled using high-level functions to reduce programming time. The combination of an entirely graphical programming approach with high-level functions and integrated error-correction allows novice users to operate the Interrogator even for complex experimental protocols with time-varying perfusion rates (**Supplementary Fig. S3**). By using a web application stack, the system software allows for remote access to machines over a network through a central server computer (**Supplementary Fig. S4**). Multiple client browsers can communicate with a single Interrogator or a single client browser can communicate with multiple instruments; each instrument’s hardware components are connected to a Mac Mini driver computer. The driver computer runs JavaScript code that translates the experiments and actions from the client into a set of machine motions to be run on each subcomponent sequentially and/or concurrently. The driver computer also provides constant updates to the client with machine component status and the overall machine state including liquid levels, loaded consumables, and current running experiment step. Networking the system platform enables remote experimental design, operation monitoring, and deployment of software updates.

Using the microscope module of the Interrogator (Fig. 1e), still images and dynamic recording of organ actuation are equally possible, with the ability to visualize, for example, the villi-like morphology and the cyclic stretching of the Gut Chip by applying cyclic suction to the hollow side microchambers (Fig. 1f, **Supplementary Fig. S1, Supplementary Movie 3**). The challenge of providing phase contrast imaging, which is required for non-invasive visualization of some tissues, was solved using a phase plate suspended between the Organ Chip cartridges and an LED array light source (**Supplementary Fig. S5**). Due to the modular design, the alignment of the Organ Chips relative to the rectangular slits of the phase plate serve to structure light from an LED array to create a partial phase contrast light source. The microscope controls allow regions of interest (ROIs) to be saved and returned to for time-lapse imaging, and to facilitate repeated inspection of the same ROIs over time in the Organ Chips.

### Interrogator characterization

An instrument calibration procedure was developed and validated by measuring the positional error of the system prior to and after calibration, which greatly improved positional accuracy (Fig. 2a,b). Prior to calibration, the instrument had positional errors –4.07 to +1.70 mm across the three axes due to inherent part variation, misalignment during assembly, and deflection of the axes during operation (Fig. 2c). As expected, the positional errors for each axis were dependent on the position relative to the other axes (e.g., the positional error of the z-axis was dependent on the position of the x-axis). Importantly, after calibration, the positional error for all three axes remained less than 0.5 mm (Fig. 2d). The stability of each axis was tested using an automated routine of 1,000 cycles of motion through the full range of motion; success corresponded to maintaining positional stability to within 0.1 mm, which was accomplished throughout all cycles. The pipettor also was characterized for accuracy and precision of fluid handling using an automated serial dilution protocol as well as repeated pipetting of volumes. The standard dilution data were fit to a four-parameter logistic regression (4PL) with a correlation coefficient R^2^ = 0.9998 (Fig. 2e). Pipetting error was less than 2% for volumes of 100 µL and above when using 1 mL pipette tips, and only 4 % for a volume of 50 µL at the extreme low volume capability range (Fig. 2f).

**Figure 2.**
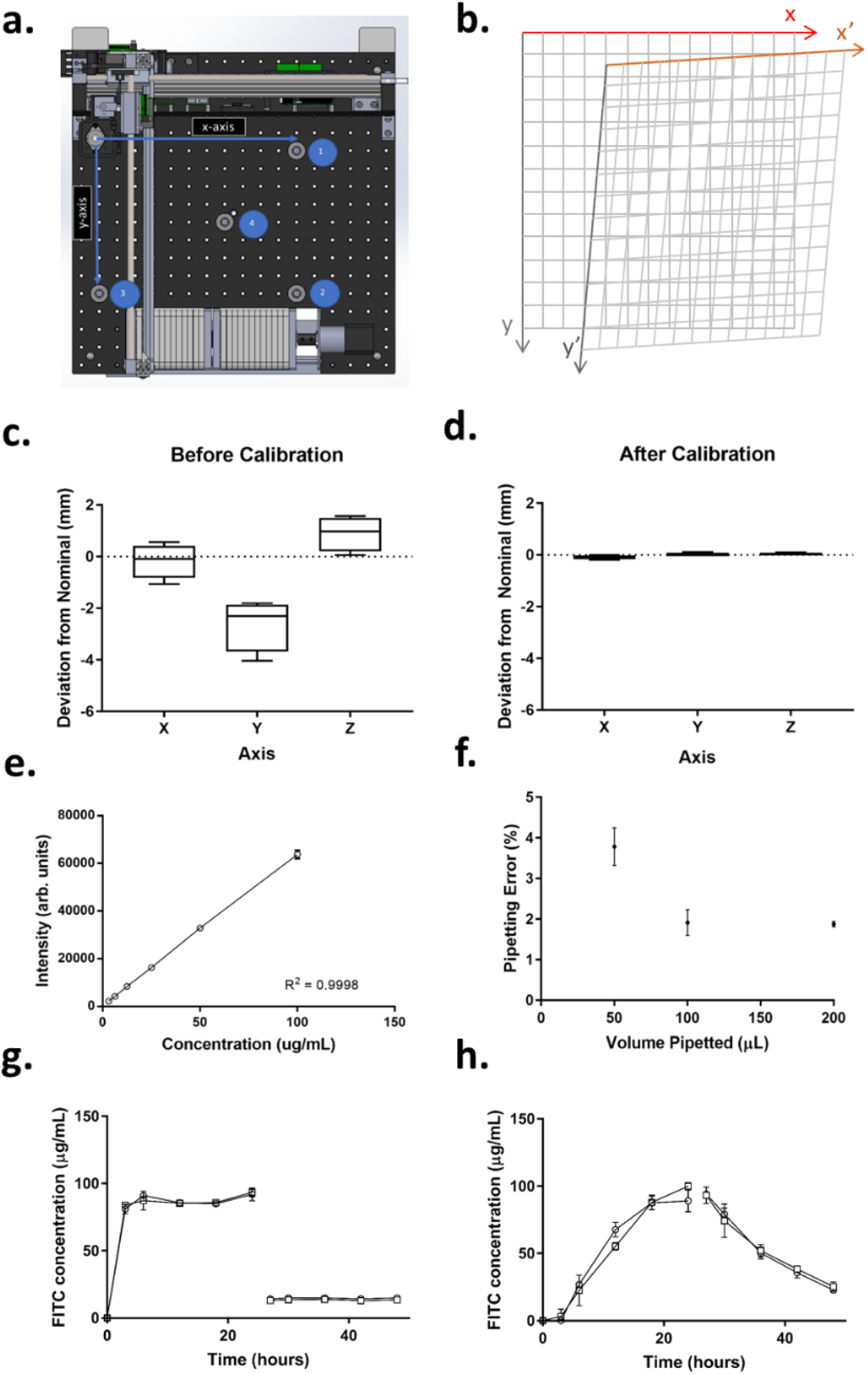
Schematic of stage calibration. **(a)** A matrix of x,y,z coordinates is corrected by touching the four calibration points with a probe attached to the pipettor head. **(b)** The actual coordinates are then mapped to the theoretical coordinate matrix to generate a calibration matrix that is automatically used to adjust fluid handler stage motion. Initial positional errors **(c)** can be reduced below 0.5 mm **(d)**. **(e)** Accuracy and precision of liquid handler used in Organ Chip linking studies was measured using a standard dilution routine of fluorescent dye. N=8. **(f)** Accuracy of pipettor calculated as Error = SD/Mean. Error bars are SE of error and correspond to the precision. N = 60. Automated generation of absorption/adsorption calibration data using blank Organ Chips and inulin-FITC tracer dye during infusion and elution phases. The inlet reservoirs **(g)** show the addition of fluorescent tracer for the first 24 h, while the outlet reservoirs **(h)** exhibit dye dilution due to perfusion through the Organ Chip device.

The modularity of the system lends itself to numerous design configurations. For example, to develop model calibration parameters for an inulin-FITC tracer dye that is commonly used to characterize tissue permeability barriers *in vivo*^36–38^, as well as in Organ Chips^24,29,39^, the Interrogator was used to determine tracer distribution, as well as its absorption and adsorption by the PDMS chips and associated materials under conditions in which perfusion was carried out using empty Organ Chips without cells. The Interrogator automatically executed the tracer dosing and elution study by pipetting a solution containing the fluorescent tracer into the apical and basal inlets of poreless membrane chips, perfusing medium through the apical and basal channels of the Organ Chips using the integrated peristaltic pump, and performing robotic sampling of their outlets over 24 h. This was followed by a 24 h washout phase where the pipettor introduced fresh medium into the inlets without tracer dye. Inlet fluorescence rapidly reached steady state during the infusion phase (Fig. 2g), while a gradual increase in fluorescence intensity was seen in the outflow samples (Fig. 2h). In addition, desorption of fluorescent tracer was observed during washout phase (Fig. 2h), and these measurements were then used in the quantitative analysis of a HuBoC platform containing 8 different human Organ Chips fluidically linked using the Interrogator Instrument.

### Leveraging the Interrogator to create an automated HuBoC platform

Given its capabilities, the Interrogator instrument should enable a range of studies from automated culture of single Organ Chips to coupling of chips for first pass metabolism to more complex whole HuBoC analysis. To create a HuBoC platform, we used the Interrogator to fluidically link, perfuse, and culture 8 different human Organ Chips representing: intestine, liver, kidney, lung, heart, skin, BBB, and brain that were lined by human parenchymal cells from each these organs in one channel and vascular endothelium in the parallel channel (Fig. 3a,b). The methods for creating each of these chips have either been published^13,31,34,35,40^ or are described in **Methods**; note that to increase the absorptive surface area of the Gut Chip, the channels were lengthened relative to past publications by creating a serpentine pattern on-chip (Fig. 3b). Each of the Organ Chips was cultured until it reached a mature state, as determined by measuring organ-specific functions (i.e., intestinal villi formation and barrier function; liver albumin production; kidney albumin reabsorption; heart contractility; lung barrier function; skin cell differentiation and barrier function; BBB permeability function **(Supplementary Fig. S6)**, and the time required differed depending on the organ type **(Supplementary Fig. S7**).

**Figure 3.**
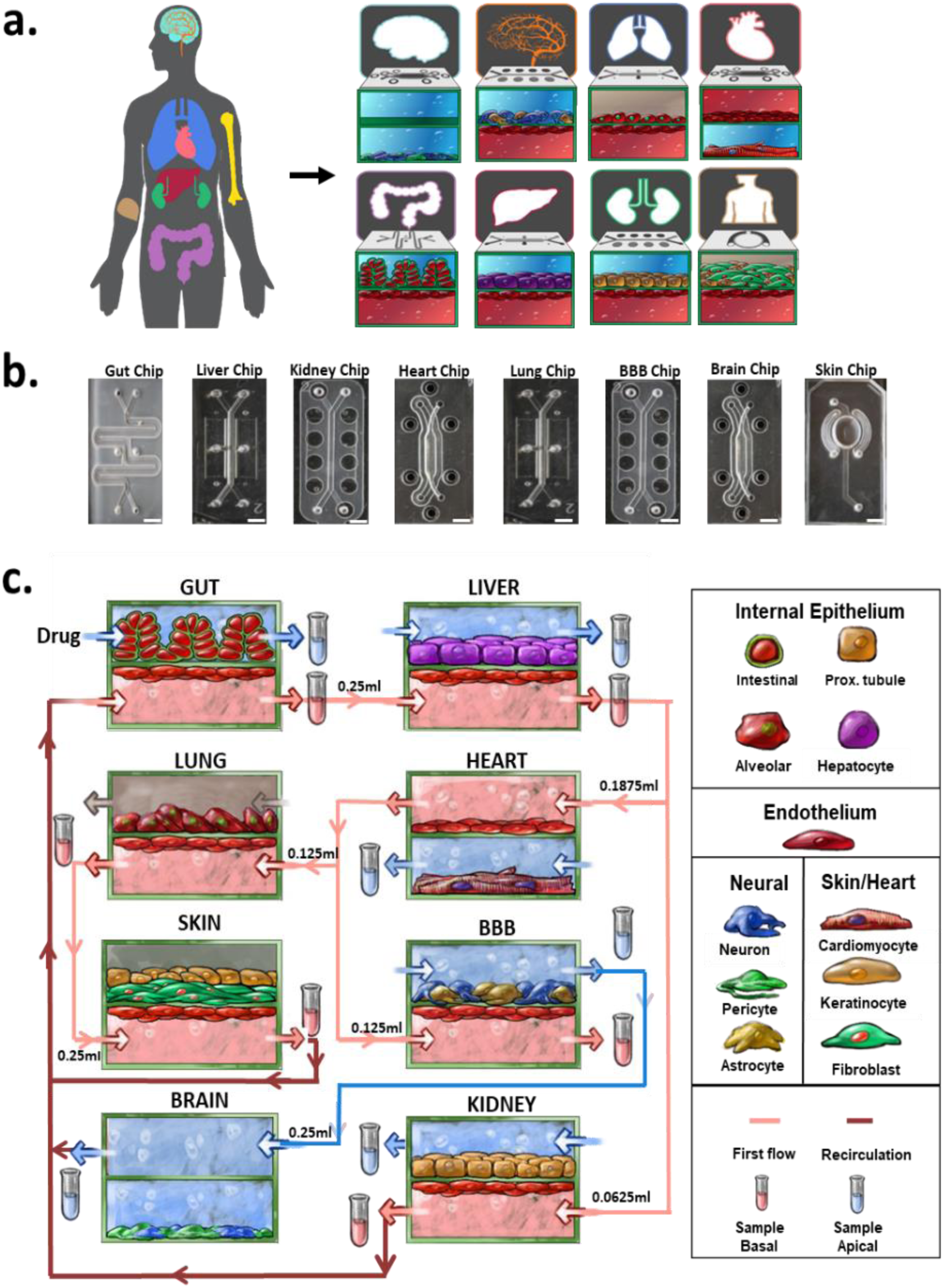
Linking scheme of 8 organ Human Body-on Chips. **(a)** Human organs for the Human-Body-on-Chip (HuBoC), each colored organ has a developed representative Organ Chip. Upper row, left to right, Brain: differentiated human primary neural stem cells forming networks of neurons and astrocytes. BBB: lower channel brain microvascular endothelial cells, upper channel brain pericytes and cortical astrocytes. Lung: lower channel human umbilical cord vascular endothelial cells, upper channel, lung epithelial cells. Heart: lower channel human umbilical cord vascular endothelial cells, upper channel cardiomyocytes differentiated from cardiomyocytes. Gut: lower channel human umbilical cord vascular endothelial cells, upper channel containing villi-like structures of gut epithelial cells, Liver: lower channel containing liver sinusoidal endothelial cells, upper channel containing hepatocytes. Kidney: lower channel kidney derived endothelial cells, upper channel containing proximal tubule epithelial cells. Skin: lower channel containing dermal microvascular endothelial cells, upper channel containing keratinocytes and dermal fibroblasts. **(b)** Photographs of the Organ-Chips: Gut Chip, Liver Chip, Kidney Chip, Heart Chip, Lung Chip, BBB Chip, Brain Chip, and Skin Chip. Scale bar: 5mm. **(c)** A total of 8 vital organs – Gut, Liver, Heart, Kidney, Lung, Heart, Brain, Blood Brain Barrier (BBB)), and Skin – were joined through vascular endothelial channels in order to create the Body-on Chips. The system enables multiple sampling points and variety of linking possibilities. The numbers represent the transfer volumes per linking step.

These Organ Chips were fluidically linked to mimic oral dosing of a compound through the lumen of the Intestine Chip and subsequent distribution to the body. The apical channels were perfused at the same rate as the basal vascular channels, and fluidic linking was carried out every 12 h by transferring small volumes (62.5-250 μL) of blood substitute culture medium from the outflows of the Organ Chip vascular channels to the inlets of the other linked Organ Chips in the order described in Fig. 3c. This linked fluidic network was maintained at a 1 μL/min flow rate; periodic 5 μL/min flush cycles were also applied automatically using the integrated peristaltic pump and control software to dislodge any debris or bubbles that could impede perfusion during the 3 weeks of culture. Because the vascular channels of the Organ Chips were lined by a continuous endothelium, it was possible to perfuse multiple different linked Organ Chips with a single endothelial medium – the ‘blood substitute’ – to mimic *in vivo* blood perfusion of multiple organ systems.

The automated Interrogator instrument effectively maintained perfusion, viability, morphology, and organ-specific functions of all 8 Organ Chips throughout the entire 3 weeks of fluidically-linked perfusion culture in two separate studies (Fig. 4). The responses of the HuBoC Organ Chips were analyzed with multiple organ-specific structural and functional assessments including immunostaining for VE-Cadherin to measure endothelial integrity on all Chips (except the Brain Chip that did not have an endothelium as it was linked to the BBB Chip) and for ZO-1 to assess epithelial integrity in the Gut, Lung and Kidney Chips; apparent permeability (P_app_) values for Inulin-FITC, dextran (3 kDa), or Cascade blue (596 Da) were also used to estimate barrier function in the Gut, Lung and Skin Chips, respectively; MRP2 staining of hepatocytes as well as albumin production were used to assess the Liver Chip; Kidney Chip function was measured by quantifying albumin reabsorption; α-actinin staining and LDH release were used to analyze cardiomyocytes in the Heart Chip; epidermal cells were stained with loricrin and keratin 14 in the Skin Chip; GFAP staining and LDH release were used to measure function and viability of astrocytes and pericytes in the BBB Chip; and Brain Chip function was assessed by staining for astrocytes (GFAP) and neurons (β-III-Tubulin) as well as by quantifying the glutamine:glutamate ratio via mass spectrometry. Immunofluorescence staining of these various cell markers confirmed maintenance of tissue-specific morphology and relevant distributions of molecular markers in all Organ Chips (Fig. 4, **top**), confirming low cell death rates.

**Figure 4.**
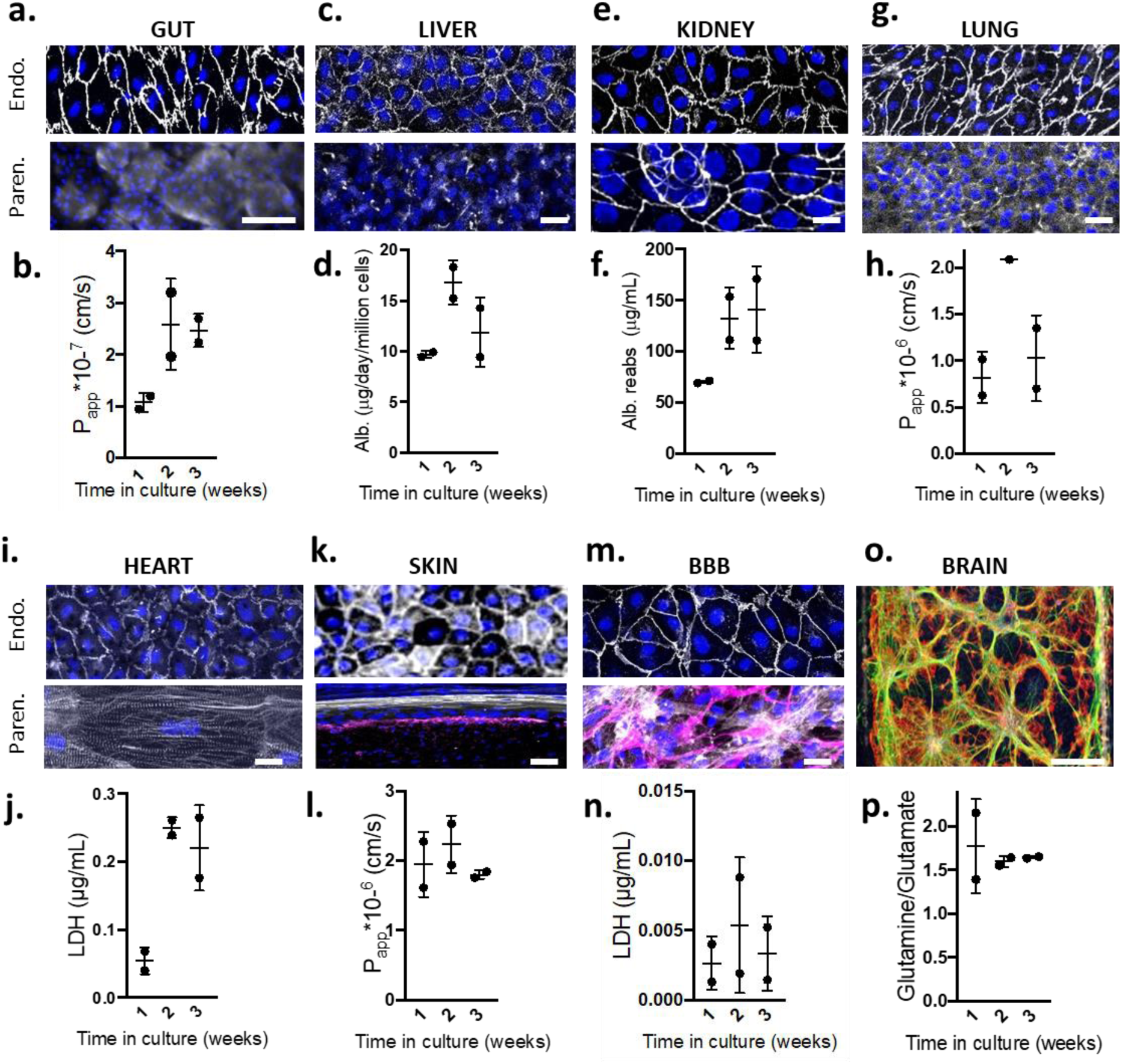
Automated Human Body on Chips linkage demonstrates maintenance of organ viability and function for 3 weeks. Immunofluorecence imaging of the HuBoC organs and organ-specific assessment throughout 3-week linkage: **(a)** Gut Chip endothelium (VE-Cadherin) and parenchyma (ZO1); **(b)** Gut Chip permeability values for Inulin-FITC; **(c)** Liver Chip endothelium (VE-Cadherin) and parenchyma (MRP2); **(d)** Liver Chip albumin production; **(e)** Kidney Chip endothelium (VE-Cadherin) and parenchyma (ZO-1); **(f)** Kidney Chip albumin reabsorption; **(g)** Lung Chip endothelium (VE-Cadherin) and parenchyma (ZO1); **(h)** Lung Chip dextran (3 kDa) permeability; **(i)** Heart Chip endothelium (VE-Cadherin) and parenchyma (α-Actinin); **(j)** Heart Chip LDH secretion; **(k)** Skin Chip endothelium (VE-Cadherin) and parenchyma (white Loricrin and purple Keratin 14); **(l)** Skin Chip cascade blue^®^ (596 Da) permeability; **(m)** BBB Chip endothelium (VE-Cadherin) and parenchyma (white pericytes, purple astrocytes (GFAP)); **(n)** BBB Chip LDH secretion; **(o)** Brain Chip parenchyma (green astrocytes (GFAP) and red neurons (β-III-Tubulin)); **(p)** Brain Chip Glutamine:Glutamate ratio. Scale bars: 100 µm. Data recorded at the given time points from two independent experiments; micrographs acquired from unlinked control chips following 3 weeks of culture. The IHC is representative of multiple Organ Chips and Chip regions.

To verify control of medium perfusion throughout the entire HuBoC *in vitro* model and quantify distribution of soluble small molecules, we infused inulin-FITC tracer dye ‘intravenously’ into the vascular channel of the Gut Chip once every week to mimic the distribution of a small molecule or drug immediately after it has been absorbed through the intestine lumen into the vasculature. The tracer dye concentrations were highly reproducible in all Organ Chips when measured over the 3-week period of culture (Fig. 5). This result confirms the stability of the fluidically linked HuBoC platform and all levels of Interrogator operation, from perfusion rates to pipettor distribution and sampling accuracy to maintenance of Organ Chip barrier function over 3 weeks of linking.

**Figure 5.**
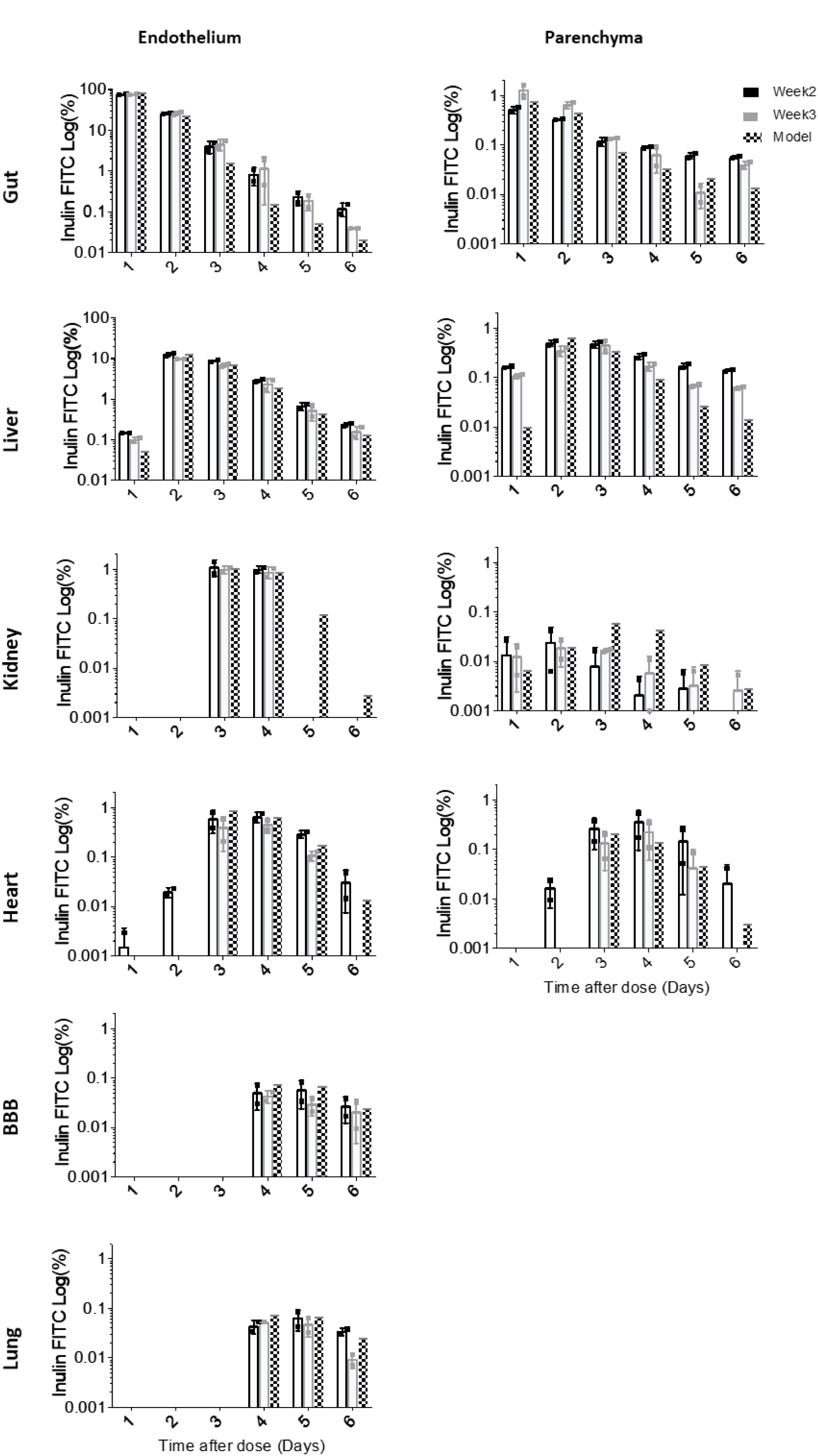
Long-term analysis of inulin-FITC (2-5 kDa) tracer dye pharmacokinetics in an 8 organ system linked via vasculature and supported by computational PBPK modeling. Inulin-FITC was given as a bolus dose once weekly on the parenchymal side of the Gut Chip and linked as shown in Fig. 4. Experimental values (white bars, days after dose of the second week and gray bars, days after dose of the third week) for Inulin-FITC concentration throughout the linked Organ Chips match with the PK model predictions (black bars) over 21 days in both the endothelial and parenchymal channels. Data from two independent experiments. Read-outs below 0.005% are at the limit of detection.

To further understand whether the distribution of inulin-FITC tracer within the linked Organ Chips of the HuBoC platform was quantitatively relevant (i.e., in addition to being stable and reproducible), we developed a physiologically-based MCRO computational model of the linked Organ Chips derived from first principles. First, detailed 3D simulations of the Organ Chips were created, incorporating convection and diffusion mass transport, physicochemical material properties of absorption and adsorption, for each of the specific Organ Chip and fluidic linkage geometries (**Supplementary Fig. S8)**. As the same chip designs were used for the Gut, Kidney, Liver, and BBB Chips, we used the standard Organ Chip model to simulate these chips; specialized models were created for the Lung, Skin, and Heart/Brain Chips (**Supplementary Figs. S9-11**).

The model was calibrated for inulin-FITC absorption and adsorption kinetics using data shown in Fig. 2g,h. To accelerate the simulation time, we converted the computationally intensive 3D computational organ models into reduced order systems of ordinary differential equations. Using this approach, we developed fast-running MCRO models for all planar Organ Chip designs except the Heart Chip, where a 3D model was used to capture the motion of the cantilevered muscular (cardiomyocyte) thin films that are included in the parenchymal channel of this chip (**Supplementary Fig. S11**). These reduced-order models were treated as convective-diffusive plug flow reactors rather than the commonly-used well-stirred reactors, which more accurately mimics *in vivo* organ perfusion and provides meaningful residence time information. This *in vitro* MCRO model solves convection-diffusion-partition-reaction equations for species in the perfusing media, cellular barriers, in membranes, and in the PDMS package material. Importantly, this MCRO model recapitulated the experimental concentrations of inulin-FITC over time within all 8 Organ Chips with good agreement (Fig. 5), confirming that the Interrogator instrument enabled robust quantitative experimentation in multiple linked Organ Chips within this HuBoC platform. Analysis of organ-specific assays at the conclusion of the tracer study, once again confirmed that the functions of all 8 Organ Chips were maintained over 3 weeks even when subjected to this automated experimental protocol (Fig. 3).

## DISCUSSION

The ability of animal models to quantitatively predict drug toxicity and efficacy in humans is limited, which has prompted significant efforts focusing on developing human *in vitro* models for PK/PD testing^1,6,41,42^. Vascularized Organ Chips have been shown to recapitulate human organ-level physiology as their two-compartment, semi-permeable membrane structure mimics *in vivo* communication and mass transport across physiologically relevant tissue-tissue interfaces with the endothelium-lined vasculature^1,2^. In addition to mimicking healthy organ physiology, Organ Chips have recapitulated pathophysiological conditions, including chronic-obstructive pulmonary disease, tobacco smoke exposure, asthma, and viral infections in the small airway^43,44^, bacterial infections^34,45^ and radiation exposure^46^ in the gut, and drug toxicities in multiple Organ Chips^31,39,47^. While these approaches highlight the value of single Organ Chips, PK/PD analysis requires multi-organ or whole-body systems linked by vascular perfusion.

Here we describe an automated Interrogator instrument that can perfuse and fluidically link up to 10 different human Organ Chips and maintain their viability and function for at least 3 weeks in culture using a common blood substitute medium. Although robotic fluid handlers have been previously used for microfluidic tissue culture, the Interrogator represents the first system capable of complex, programmable experiments using perfused single or linked Organ Chips. The robotic Interrogator instrument supported Organ Chip culture through integrated perfusion, automated replenishment of media, precision fluid transfers between linked Organ Chips, and sampling of Organ Chip inlets and outlets for analysis of inulin-FITC distribution. We also demonstrated that the Interrogator can be leveraged to create a viable HuBoC platform that maintains multi-organ function for at least 3 weeks *in vitro*. Although only a branched serial linkage scheme with 8 Organ Chips was used here, robotics-based organ linking can be readily reconfigured to include varying numbers of organs as well as an arterio-venous reservoir^48^. The modular design of both the Interrogator hardware and control software enables rapid addition or removal of Organ Chips without disrupting the experiment, including, importantly, the perfusion of media. The dosing of drugs can be modified to any mode of delivery: oral dosing to the gut, intravenous dosing to the interconnected vascular medium, topical skin dosing, and aerosol dosing to the lung are all feasible to integrate. The modularity of our HuBoC approach also facilitates time-delayed linking of organs (e.g., to mimic temporary loss of organ function or perfusion) and isolation of organ subsystems, which would be difficult or impossible in animal models.

The Interrogator facilitated experimental design and model calibration for complex studies of linked Organ Chips within a HuBoC platform. Error checking integrated into the control software prevented mistakes in the fluid handling programming by continually simulating the input values to assess physical feasibility, accounting for Organ Chip perfusion over time, and volume changes due to sampling, linking, and media replenishment. Furthermore, the software alerted users if inadequate pipette tips or media volumes were present. The precision of data obtained from Interrogator-automated collection of time-resolved samples during blank Organ Chip perfusion and complex linking of arbitrary Organ Chips facilitated calibration of MCRO computational models of the experimental design. This in turn enabled analysis of experimental data, which would otherwise be difficult to validate, and showed *in vitro* and *in silico* agreement. This approach can be further exploited for optimization of experimental conditions *in silico* by enabling simulation of tracer distribution through the linked Organ Chips to assess optimal sampling time points and study duration. The capabilities enabled by the Interrogator demonstrate how close interplay between computational and experimental efforts may accelerate PK/PD experimental design and data analysis in HuBoC platforms^7^.

PBPK models have relied on conventional *in vitro* cell culture devices, such as Transwells, to determine specific input parameters, such as drug solubility, barrier function, drug partition coefficients, intrinsic clearance rate (CL_int_) in the liver, and renal clearance rate in kidney^49,50^. However, these static, quasi-equilibrium, *in vitro* platforms cannot replicate key dynamic physiological organ characteristics: micro-organization of tissues within organs, perfusion of blood and other bodily fluids that impact drug metabolism and transport, time-dependent dosing of drugs and metabolites that cross back and forth across the endothelial-tissue interface, organ-organ interactions, and chemical, biological, and physical microenvironments specific to the respective tissues. Mathematical *in vitro*-to-*in vivo* extrapolation (IVIVE) models have been developed to address some of these limitations, but these often fall short due to non-physiological scaling factors and organ structure assumptions (e.g., the liver is viewed as a well-stirred reactor)^51,52^.

Microfluidic Organ Chip devices with perfused endothelial and parenchymal channels and multi-organ microphysiological systems have the potential to overcome these limitations^6,7,9,53^. For example, unlike most past PK models, here we leveraged the biomimetic design of Organ Chips to conduct full 3D simulations of mass transport without the pitfalls of well-stirred reactor assumptions. Incorporating the fundamental equations of previous PBPK models^51,54^ and adapting them to describe a more physiologically accurate *in vitro* experimental system resulted in the MCRO model described here. Importantly, the equations we derived are also equally applicable to analyze human organ mass transport *in vivo* to better model human physiology^53^. This model and simplified reduced-order differential equation models were used to confirm the quantitative accuracy of the Interrogator-enabled HuBoC platform using an inert inulin-FITC tracer dye in the present study, which is focused on validation of the Interrogator instrument. However, in a parallel effort, this model has been extended to create an MCRO model that incorporates drug transport and metabolism with subsequent *in vitro-in vivo* translation^48^.

While others have demonstrated fluidic linking of multiple microfluidic devices, they were each linked by a single type of organ parenchymal cell (e.g., liver, lung or kidney epithelium)^10,55^, and thus, their lack of a separate vascular compartment with associated flux between the parenchymal and vascular spaces limits their use in PBPK models and extrapolation to humans. Furthermore, the direct contact of parenchymal tissues with a shared medium not only poses a significant technical challenge for developing a universal blood substitute that all cell types can tolerate, but it may hinder normal tissue function by forcing parenchymal cells of different organs to interact directly, which does not occur *in vivo*. Several microphysiological systems have incorporated tissue culture well inserts with semi-permeable membranes lined by both parenchymal cells and endothelial cells to develop a more physiological shared medium for linked organs^11,19,20^. Yet others have demonstrated 3D culture of endothelial cells with a parenchymal tissue in single organ configurations^56^, which facilitates a shared medium but presents additional variability in vasculature area for PBPK calculations. Despite these efforts to incorporate a shared vasculature, the lack of perfused parenchymal compartments in these systems still hinders PK/PD modeling as it does not incorporate the *in vivo*-like convection on the parenchymal tissues that is present in nearly all organs (e.g., urine flow, bile flow, cerebrospinal fluid flow).

The Interrogator-enabled HuBoC platform approach presented here addresses all of these problems simultaneously: 1) Organ Chips have parenchymal channels perfused with a relevant interstitial fluid that are interfaced across a semi-permeable membrane and with an endothelium-lined channel that is perfused by a shared universal blood substitute medium; 2) fluidic linking can be arbitrarily programmed, before or during an experiment, and can entail linking of parenchymal media (e.g., BBB parenchyma to neuronal compartment in the Brain Chip, as demonstrated recently^13^); and 3) the HuBoC linkage is amenable to first-principles-based *in silico* modeling using equations that can apply to human physiology *in vivo* as well as to Organ Chips *in vitro*. The ability to use a universal medium across Organ Chips dramatically increases experimental flexibility by leveraging the Interrogator platform for HuBoC linking with the potential to arbitrarily reconfigure the linkage scheme without the need to individually assess the viability of each new Organ Chip linkage. Importantly, we have recently confirmed the utility of this versatile Interrogator-enabled HuBoC approach coupled with MCRO modeling for evaluating drug PK/PD in a first-pass Organ Chip system and were able to recapitulate clinical drug profiles and PK parameters for two drugs (nicotine and cisplatin) previously observed in humans^48^.

There is a growing need to automate many biological *in vitro* experiments for improved accuracy, increased throughput, and reduced risk to researchers when working with pathogens or other laboratory hazards. The compact Interrogator system is a versatile tool that can be used for time-resolved studies in biocontainment scenarios where human experimenters could be placed at risk, such as when studying the pathogenesis of human viral and bacterial infections or developing chemical weapon countermeasures. The robotic platform offers a modular, fully programmable design for Organ Chip experimentation, whether using individual multi-channel or single channel chips, or a more complex HuBoC platform. This *in vitro* human experimentation platform, coupled with physiologically-based MCRO modeling, may accelerate development of more effective therapeutics and medical countermeasures, reduce potential for drug toxicity, and optimize design of drug dosing regimens.

## METHODS

A Solidworks CAD package of the Interrogator that defines all hardware components and assembly instructions are contained in the **Supplementary Materials**. Control software is available at https://gitlab.com/wyss-microengineering/hydra-controller, and video tutorials are available at: https://vimeo.com/album/5703210

### Robotic Fluid Handling Platform

The Interrogator is mounted on a 450 mm x 450 mm aluminum optical board base (Thorlabs, Inc.) to allow for insertion into most standard tissue culture incubators. The platform consists of a pipettor (Z-Series Pipette Pump 0949, TriContinent Inc., Grass Valley, CA, USA) and control board (M-Series Controller PCBA 0955, Tricontinent Inc., Grass Valley, CA, USA) mounted on a custom three-axis gantry. The motion system consists of X-Y stepper motors (Adafruit #324) paired with a 14-1/2 degree pressure angle 32-pitch rack and 16-tooth pinion system. The Z-axis is comprised of a stepper motor and screw drive (RoboDigg 11HY0401-200T52). Limit switches are used on the proximal end of all three axes, with a 1mm homing offset. The gantry is controlled via g-code commands sent to a GRBL Arduino shield (settings described in **Supplementary Fig. S12**). Perfusion is achieved using two stacked 12-channel peristaltic pump heads (DG-12-B/D, Longer Pump Inc., Boonton NJ, USA) controlled by a geared stepper motor and custom control via an Arduino. Individual Organ Chips are mounted in stainless steel cartridges that each hold two inlet and two outlet reservoirs. Cartridges are snapped into a spring-loaded carrier tray and tubing is connected to the peristaltic pump. Two standard well plates, two pipette tip boxes, and a waste collection bag are mounted in the rest of the platform deck.

### Platform Validation

The gantry was optimized and validated by speed testing until failure to establish maximum speed set points, motion stability testing to within 0.1 mm over 1,000 motion cycles, and positioning accuracy and precision across the platform by puncturing 96 well plates using a pipette tip. The peristaltic pump was validated for perfusion stability and accuracy using a Sensirion SLI-0430 flow meter over 72 hours. Pipettor accuracy was assessed using automated dilution of fluorescein dye, and precision was measured by 48 repeated transfers of fluorescein dye at 100 and 200 µL volumes.

### Microscope Module

The microscope module was designed in SolidWorks (Dassault Systèmes; see **Supplementary Materials** for CAD files) and shown in Fig. 1e. The three-axis positioning stages were machined from corrosion-resistant aluminum 6061. Acme threaded rods were coupled to pancake stepper motors (44M100D, Portescap, West Chester, PA) for compact actuation capability, and dual rails with ball bearing carriages were used for smooth translation movement of the separate axis assemblies. The motors were driven via a GRBL shield mounted on an Arduino Uno controlled through Python. The mechanical components of the microscope module were bolted to the optical breadboard base of the fluid handling robot.

The optical subassembly of the microscope consists of 1-inch diameter optical tubes (Thorlabs SM1 series) connected to a mirror cube and Nikon objective adapter ring. In this study, a 10X Nikon phase contrast objective was used. Images were acquired via a Python script from a camera (Basler cA2500-14um). The image was focused using a 50 mm focal length achromatic doublet lens (Thorlabs AC254-050) via a 100 mm focal length lens to reduce optical path distance. To provide appropriate illumination for observing delicate cell features across the imaging area without a condenser, a phase plate was designed to be interchanged with the well plate holders. The phase plate **(Supplementary Fig. S5)** consisted of a black acrylic sheet, laser cut with slits that correspond to the objective phase ring diameter. Illumination for the microscope assembly was provided by a strip of white LEDs mounted to the top inner surface of the incubator.

### Software Architecture

A JavaScript web interface communicates with the platform hardware *via* a server, enabling remote control and troubleshooting access. The web interface facilitates experimental protocol setup using a completely graphical user interface. Users can set up reservoir and well plate locations, program fluid transfer time points, and control the perfusion system in an integrated display. In addition, the user interface contains real-time volume calculation during the experimental design step that incorporates volume changes due to perfusion and fluid transfers, offering a layer of error correction to accelerate experimental program validation.

The web application uses a variation of the MEAN stack application. The MEAN stack is comprised of NodeJS and ExpressJS as the server, MongoDB as the database, and AngularJS as the front-end model-view-controller framework **(Fig. S4)**. The benefit of the MEAN stack is that it is written completely in Javascript, requiring less cross-language data parsing and developer knowledge overhead. The control software uses a specific construction of the MEAN stack, called Angular Fullstack with the addition of SocketIO, for real time socket communication between client, server, and machine. Numerous tutorials on the MEAN stack and SocketIO can be found online.

### Liquid Handler Characterization

Precision of the liquid handling system was characterized by programming the 3-axis robotic pipettor to create a serial 2X dilution using inulin-FITC dye and then measuring fluorescence of solutions using a BioTek Synergy Neo plate reader with 485 nm excitation. Eight replicates of the dilution series were pipetted to allow statistical analysis. Thorough mixing was performed before and after each transfer.

Reproducibility was measured by pipetting aliquots of 100 µg/mL inulin-FITC solution from a supply reservoir to a 96 well plate and then measuring fluorescence using a plate reader as above. Aliquots of 50, 100, and 200 µL were pipetted in two batches of approximately 30 samples per condition in two separate experiments. The mean and standard deviation of each volume within an experiment were used to calculate accuracy; the mean and standard deviation between experiments were used to calculate reproducibility.

### Microfluidic Device Fabrication

The gut, liver, lung, kidney, skin, and BBB chips were fabricated as previously described^23,24,57^. Briefly, molds for the microfluidic devices were fabricated out of Prototherm 12120 using stereolithography (Protolabs, Maple Plain, MN). The top and bottom components of the devices were cast from polydimethyl siloxane (PDMS) at a 10:1 w/w base to curing agent ratio and bonded to a porous PDMS membrane using oxygen plasma (Atto, 30 mbar O2, 50 W, 2 min; Diener Electronic GmbH, Ebhausen, Germany). The membranes provide a semi-permeable barrier between the epithelium and microvascular endothelium layers and were fabricated by casting against a DRIE-patterned silicon wafer (50 × 50 mm) consisting of 50 μm high, 7 μm diameter posts spaced 40 μm apart. Chips with PDMS membranes enable application of cyclic strain via programmable sinusoidal vacuum pressure applied to the vacuum ports parallel to the fluidic channels. Chips fabricated with polyester terephthalate (PET) track-etched membranes were assembled using (3-Glycidyloxypropyl) trimethoxysilane (GPTMS; Sigma, 440167) and (3-Aminopropyl) triethoxysilane (APTES; Sigma, 440140) as previously described^57^. Detailed chip dimensions are described in **Supplementary Table 1** and chip cross sections are show in **Supplementary Fig. S1**.

#### Heart Chip

This chip is composed of two parts of polycarbonate and a porous membrane. The two PC parts were designed with SolidWorks software (SolidWorks Corp., Waltham, MA, USA) and produced by micromachining. The PC parts were then sonicated twice for 15 min in soapy water and once in water to remove residual oils from the machining process. Thereafter the PC parts were dried with condensed air and incubated overnight at 65 °C for drying. In order to polish the surfaces, the PC parts were briefly exposed to dichloromethane, DCM (Sigma-Aldrich, St Louis, MO, USA), vapors and dried in a dust-free environment at room temperature for 24 h. The PC porous membrane was cut with a UV laser (Protolaser U3, LPFK Laser and Electronics, Garbsen, DEU). The chip was placed in a vacuum chamber containing 4 mL of DCM for 30 min to allow bonding of the parts. The PC porous membrane was sandwiched between the two PC parts, manually aligned, and compressed at 130-140 °C and 150-200 psi for 8 h. After the bonding treatment, the chips were ready to use. A chip base and manifolds were designed with SolidWorks (SolidWorks Corp., Waltham, MA, USA) and fabricated in PC by micromachining.

#### Brain Chip

Uses same procedure as Heart Chip, but with the following additional steps: After 2 weeks of neural cell culture, the TOPAS^®^ (PolyLinks, Asheville, NC, USA) substrates were assembled into the chip. The TOPAS^®^ was first deposited in the bottom of the base. Then, a PDMS gasket (molded with an aperture corresponding to the neuronal growth area) was placed on top of the TOPAS^®^ substrate. The PC chip was finally placed on top of the gasket and the whole was maintained on the base by screwing the manifolds to the base. Media reservoirs were fixed on one manifold. Reservoirs consisted of 5 mL syringes from which the top was cut. The plungers were cut and a biopsy punch used to create a minimal opening to the atmosphere. Connectors were fixed to the other manifold. The connector linked to the bottom channel (neuronal channel) was blocked and the connector linked to the upper channel was connected to a peristaltic pump (IPC series 16 channels, Ismatec, Cole-Parmer, Wertheim, DEU). This configuration prevented any shear stress on the neuronal constructs while enabling diffusion through the PC porous membrane.

#### Skin Chip

The skin chip design reflects the requirement for a thicker tissue layer that needs to be uniformly strained. To accomplish this, the skin apical channel is a static oval reservoir with vertical posts around the perimeter. The posts enable collagen to gel around them, providing mechanical support for applying strain. The apical reservoir is open at the top to allow for deposition of viscous collagen and cells and is resealable with a medical grade adhesive film (Adhesives Research).

#### Mechanical actuation of chips

Microfluidic gut, lung, and skin cultures were mechanically actuated using a programmable vacuum regulator system built in-house. The system consists of a vacuum regulator (ITV0091-2BL, SMC Corporation of America, Noblesville, IN) electronically-controlled by an Arduino Leonardo and MAX517 digital to analog converter. The regulator outputs a sinusoidal vacuum profile with a user-settable amplitude and frequency.

### Organ Chip Culture

Organ Chips were sterilized using oxygen plasma prior to applying an optimized extracellular matrix coating and subsequent seeding of epithelial and endothelial cells. Cell culture parameters are detailed in **Supplementary Table 2.** The chips were cultured until they reached a mature state specific to each organ type **(Figs. S6 and S7)**. Gut^48^, Liver^48^, BBB^13^, Brain^13^, Heart^40^, and Kidney^48^ Chip cultures were performed as previously described. The Lung Chip was prepared as published^35^, except A549 alveolar epithelial cells (ATCC CCL-185) and human umbilical vein endothelial cells (ATCC PCS-100-010) were used.

#### Human Skin Chip

Prior to plating cells, the chips were surface treated with an oxygen plasma (Atto, Diener Electronic GmbH, Ebhausen, Germany) at 30 mbar, 50 W, 2 min). The basal channel was then coated with a solution of 50ug/mL human fibronectin (Corning, 354008) and 100ug/mL bovine collagen I (Gibco, A10644-01) dissolved in cold DMEM+1% Pen/Strep. Additionally, an acellular collagen solution consisting of 5mg/mL rat tail collagen was deposited on the apical membrane to help anchor the cell-laden gel to the PDMS membrane.

Human primary dermal fibroblasts derived from adult female forearm (ThermoFisher, C0135C) were cultured in DMEM-High Glucose with Glutamax (ThermoFisher, 10569) supplemented with 10% FBS. Cells were plated at 6,000 cells/cm^2^ in standard cell culture flasks and used up to P7 for all experiments. Human primary neonatal epidermal keratinocytes derived from foreskin (ATCC, PCS-200-010) cultured in keratinocyte growth medium (Dermal cell basal medium from ATCC, PCS-200-030, supplemented with keratinocyte growth kit (ATCC, PCS-200-040). Cells were plated at 13,000/cm^2^ and used up to P5 for all experiment. Human dermal microvascular endothelial cells (HDMVECs, ATCC PCS-110-010 were cultured in Vascular Cell Basal Medium (ATCC PCS-100-030) supplemented with Endothelial Cell Growth Kit-VEGF (ATCC PCS-100-041). Cells were plated at 1×10^4^ cells/cm^2^ in standard cell culture flasks and used up to P5 for all experiments.

Human adult fibroblasts were harvested and re-suspended at 1.5×10^6^ cells/mL in dermal proliferation media (ThermoFisher, DMEM-High Glucose supplemented with 1% Pen/Strep, 5% BCS, and 50ug/mL Sodium Ascorbate). Dermal collagen solution (2.5 mg/mL bovine collagen I (Gibco, A10644-01) in 1X MEM; collagen was initially pH-adjusted by mixing with 5.5% v/v 1N NaOH prior to the addition of MEM) was combined with the fibroblast suspension to achieve a final density of 3 x10^5^ cells/mL in collagen gel; 90 μL of this cell-gel suspension were pipetted into the apical reservoir of each chip and cultured for 3 days. Human primary neonatal epidermal keratinocytes (HEKn) were harvested and re-suspended at 10.5×10^6^ cells/mL in epidermal proliferation media (ThermoFisher; DMEM/F12 3:1, supplemented with 1% Pen/Strep, 0.3% chelated BCS, 50 μg/mL sodium ascorbate, 0.628 ng/mL progesterone, and 10 ng/mL hrEGF). Dermal media was aspirated from apical reservoirs and 25 μL of HEKn suspension were pipetted onto each fibroblast gel (2.6×10^5^ keratinocytes/chip). Chips were incubated at 37°C for 1 h to facilitate keratinocyte attachment; chips were perfused with epidermal proliferation media for 4 days before switching to epidermal differentiation media (ThermoFisher; DMEM/F12 3:1, 1% Pen/Strep, 0.3% BCS, 50 μg/mL sodium ascorbate, 0.628 ng/mL progesterone, 265 μg/mL CaCl_2_). Air liquid Interface (ALI) was induced following 3 days of epidermal differentiation by aspirating the apical media and replacing the basal channel media with cornification media (ThermoFisher; DMEM/F12 1x 1% Pen/Strep, 2% BCS, 50 μg/mL sodium ascorbate, and EGM-2 bullet kit (Lonza)) and maintained at ALI for three weeks.

HDMVECs were seeded on the basal side of the chip on Day 28 post initial fibroblast gel seeding. Cells were harvested and re-suspended in ATCC endothelial growth media at a concentration of 5×10^6^ cells/mL. Chip basal channels were re-coated with fibronectin/collagen solution, incubated at 37°C for 1 h, and rinsed with 30 μL of warm ATCC endothelial growth media. Skin Chips were seeded with 25 μL of the HDMVEC suspension and incubated upside-down for 2.5 h at 37°C to allow for attachment of endothelial cells to the basal side of the chip membrane. Chips were then perfused with ATCC endothelial growth media overnight before switching back to cornification media.

#### Linked Organ Chip Perfusion

Mature Organ Chips were loaded into Interrogator cartridges and connected to two inlet and two outlet reservoirs using Pharmed BPT tubing and perfused at a rate of 1 μL/min with flushing of the tubing at a rate of 5 μL/min every 4 h to maintain uniform perfusion rates during several weeks of culture.

#### Universal Endothelial Medium

Medium components were from Life Technologies (Life Technologies, Carlsbad, CA, USA) unless otherwise stated. Universal Endothelial Medium was DMEM/F12 supplemented with EGM-2 Lonza Bullet Kit (Lonza, Basel, Switzerland), 0.5% FBS, 1% Pen/Strep, and growth factors (VEGF, EGF, IGF, FGFb) according to kit instructions.

### Organ Chip Analyses

#### Albumin ELISA

Media samples from inlet and outlet reservoirs of apical and basal channels of chips were sampled automatically as part of each fluidic linking cycle. Liver Chip and Kidney Chip albumin concentrations were quantified using a human albumin ELISA quantitation kit (Bethyl Laboratories Inc., Montgomery, TX, USA) per the manufacturer’s protocol. Albumin production and reabsorption rates were normalized to total protein amount. Rates were based on the difference between the inlet and outlet concentrations and elapsed times and media volumes.

#### Barrier Function

Cascade Blue^®^ hydrazide, Trisodium Salt (Thermo Fisher Scientific Inc., Waltham, MA, USA) (for Gut Chip and Skin Chip), Dextran-Texas Red^TM^ 3 kDa (For Lung Chip), Dextran-Texas Red^TM^ 70 kDA (for BBB Chip) and Dextran-Cascade Blue^®^ 10 kDa (for BBB Chip) (Thermo Fisher Scientific Inc., Waltham, MA, USA) (for BBB Chip and Lung Chip) and Inulin-FITC (Sigma-Aldrich Corp, St. Louis, MO, USA) were used as inert tracers in the medium to quantify Organ Chip barrier function. The tracers were diluted to 100 µg/mL in apical media and flowed through the chip at 60 µL/h unless otherwise described. Media inputs and perfused medium outputs were collected for all channels and measured using a fluorescence plate reader set to the respective fluorophore excitation and emission filter sets. From these data, apparent permeability (P_app_) was calculated using the following equation^24,58^:

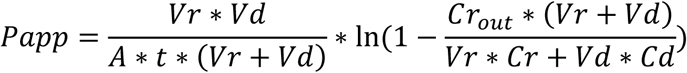

where V_R_ is defined by the volume of receiving channel effluent after time t, C_R_ is the measured concentration of tracer in receiving channel effluent, t is time of effluent collection, A is the area of the main channel, V_d_ is defined by the volume of dosing channel effluent after time t and C_d_Out_ is the concentration of dosing medium.

#### Morphology

Chips were rinsed in pre-warmed phosphate-buffered saline and fixated in 4 % paraformaldehyde (Sigma-Aldrich, St Louis, MO, USA) for 10 to 20 min at room temperature. Immunocytochemistry was carried out after permeabilization in phosphate-buffered saline with 0.05 to 0.1% Triton X-100 (Sigma-Aldrich, St Louis, MO, USA) and blocking for 30 min in 3 to 5 % Bovine Serum Albumin (Jackson ImmunoResearch, West Grove, PA, USA) or 10 % goat serum in phosphate-buffered saline with 0.05 to 0.1% Triton-X 100. Primary antibodies were applied in 2 % goat serum or 0.5 % BSA overnight at 4°C or at RT. The following primary antibodies were used for immunocytochemistry experiments: rabbit anti-glial fibrillary acidic protein (GFAP) (DAKO, 1:100, Z-0334, Lot #2002332), mouse anti-vascular endothelial (VE)-cadherin (Abcam, ab166715, Cambridge, MA, USA 1:100), (VE)-cadherin (BD Biosciences, USA 1:100, BD555661, Lot #4324666), alpha-actinin (Abcam, Cambridge, UK, 1:200), mouse anti-zona occludens-1 (ZO-1) (Invitrogen 1:100, Cat #33-9100), anti-βIII-tubulin (Sigma-Aldrich, 1:200), anti-neurofilament (Abcam, Cambridge, MA, USA 1:100), anti-glial fibrillary acidic protein (GFAP, Abcam, Cambridge, MA, USA, 1:200). Cells were washed three times in phosphate-buffered saline with 0.05 to 0.1 % Triton-X 100, followed by staining with secondary antibody staining for 30 to 60 min at RT. The secondary antibodies were anti-rabbit or anti-mouse IgG conjugated with Alexa Fluor-488, Alexa Fluor-555, or Alexa Fluor-647 (Invitrogen, Carlsbad, CA, USA). Hoechst (10 mg/mL, 33342, Life Technologies/Invitrogen, Carlsbad, CA, USA) was used at a dilution of 1:5000 for nuclei staining. For staining of F-actin, Alexa Fluor-488-phalloidin (A12379, Lot #1583098) or Alexa Fluor-647-phalloidin (Life Sciences/Invitrogen, Carlsbad, CA, USA) were used at dilution of 1:30. For glass or TOPAS^®^ bottom samples, the substrate was removed from the dish and placed on a glass slide. ProLong Gold Antifade reagent (Molecular Probes Life Technologies, Grand Island, NY, USA) was added to preserve the samples and glass coverslips were affixed using transparent nail polish. Prepared slides were either imaged immediately or stored at 4°C. Imaging was carried out in a Zeiss 710 LSM (Zeiss, Oberkochen, Germany) or Olympus confocal microscope (Olympus, Center Valley, PA, USA) with appropriate filter cubes. Image processing was done in FIJI or Imaris (Bitplane, CHE).

#### Total Protein

Lysate from the parenchymal cell channels was collected for total protein quantitation. Chips were rinsed with 1X PBS (-/-) and the basal channel was trypsinized to remove the endothelial cells. Next, RIPA Buffer supplemented with 1X Halt™ Protease and Phosphatase Inhibitor Single-Use Cocktail (Thermo Fisher Scientific Inc., Waltham, MA, USA) was added to the apical channel and incubated on ice for 20 min. Lysate was then collected from the chip and centrifuged at full speed for 10 min to remove insoluble components. Supernatant was removed and stored at −80°C until the time of analysis. Total protein was determined using Pierce BCA assay kit (Thermo Fisher Scientific Inc., Waltham, MA, USA) per the manufacturer’s protocol.

#### Mass spectrometry

All chemicals were from Sigma, St. Louis, MO, USA unless otherwise noted. To prepare media samples and calibration samples for liquid chromatograph mass spectrometry (LC-MS) analysis, 20 µL of sample was mixed with 30 µL of an internal standards solution (10 µM D4-Succinate, in Acetonitrile). After centrifugation at 18000 g for 10 minutes, 40 µL of supernatant was transferred to glass micro inserts. All samples were kept at −80°C until analysis.

LC–MS analyses were modified from Maddocks et al.^59^ and were performed on an Orbitrap Q-Exactive (Thermo Scientific, Waltham, MA, USA) in line with an Ultimate 3000 LC (Thermo Scientific, Waltham, MA, USA). The Exactive operated in the polarity-switching mode with positive voltage 3.0 kV and negative voltage 4.0 kV. Column hardware consisted of a Sequant ZIC-pHILIC column (2.1 × 150 mm, 5 μm, Millipore, Billerica, MA, USA). Flow rate was 200 μl min^−1^, buffers consisted of acetonitrile 97% in water for B, and 20 mM ammonium carbonate, 0.1% ammonium hydroxide in water for A. Gradient ran from 100% to 40% B in 20 min, then to 0% B in 10 min. After maintaining B at 0% for 5 min, it was ramped to 100% over 5 min and kept at 100% for 10 min. Metabolites were identified and quantified using Trace Finder and Compound Discoverer 2.0 software (Thermo Scientific, Waltham, MA, USA). Glutamine/glutamate standard curves were produced for quantifying those metabolites.

### Passive Tracer Distribution Analysis

Custom Organ Chips with membranes lacking pores were used to prevent any convective mixing between top and bottom channels. Chip medium was prepared by adding EGM-2 SingleQuot bullet kit (Lonza) to DMEM/F12 (Lifesciences). Inulin-FITC (Sigma Aldrich) fluorescent tracer was added to the inlet reservoirs at 100 µg/mL. 1 mL pipette tips were used for all transfers (Neptune BT1000). Fluorescence was measured in black microplates (Corning 3381 or Greiner 655096) using a BioTek Synergy Neo plate reader with 485 nm excitation. Perfusion was enabled by connecting poreless-membrane chips to the integrated peristaltic pump using 500 µm ID Pharmed BPT tubing (Thomas Scientific 1203A38) and 19 gauge stainless steel connectors (straight: Microgroup and 90° elbow: Four Slide Products). Before assembly, all chips, tubing and fittings were sterilized with oxygen plasma and to make them hydrophilic, which prevents bubble formation at junctions.

### Computational Modeling of System Behavior

All models have been developed using CFDRC’s Computational Biology (CoBi) tools^60^ (available at: http://medicalavatars.cfdrc.com/index.php/cobi-tools/). For each Organ Chip, high-fidelity simulations of coupled fluid flow, biomechanics, mass transport, biochemistry and electrophysiology models have been used to design specific organ geometry and operating parameters to reproduce the *in vivo* characteristics of the individual organs within the constraints of microfabrication and cell culture. Specifically, spatiotemporal multi-compartment reduced-order (MCRO) models of the multiple compartments were established^48^. Tissues as well as the media channels and PDMS layers were represented with a coarse spatial computational mesh represented by control volumes parallel to the media flow and one control volume in perpendicular to the direction of perfusion (PDMS, top channel, epithelial cell layer, membrane, endothelial cell layer, bottom channel, PDMS). This allowed us to solve general spatio-temporal transport equations related to accumulation, convection, diffusion, and sources of inulin-FITC and other compounds.

Computational domain of these models replicates the entire in vitro organ including microfluidic channels, PDMS membrane, epi- and endothelial cellular barriers, package material (PDMS), inlet and outlet supply tubing, and reservoirs. For the Organ Chips described in this study, a coupled perfusing media flow and conjugate mass transfer model was applied to estimate the shear stress on cell barriers, species transport and mixing, trans-barrier transport, and package material gas exchange. For organs with active mechanobiology, such as heart, gut, or lung, coupled fluid-structures interaction models were used. For example, a computational model of the Heart Chip required 3D simulation of “muscular thin film” (MTF) structures^61^ immersed in a medium and experiencing periodic mechanical contraction-relaxation movements (**Supplementary Fig. S11 and Supplementary Movie 4**). **Supplementary Fig. S11** shows the detailed geometry of multiple MTFs in the Heart Chip device, where color indicates the concentration of an adsorbed tracer compound on the MTF surface. Detailed geometry of the models used for liver, lung, and skin Organ Chips are shown in **Supplementary Fig. S9-11**. Reduced order differential equations were derived for all other Organ Chips in order to decrease computational time by integrating spatial terms for convection, diffusion, and transport into individual fluxes across control volume boundaries. Solving the convective and diffusive terms in the microchannels is achieved with second order accuracy in the direction parallel to media perfusion and analytically perpendicular to the direction of perfusion. This approach resolves spatial concentration gradients (e.g. liver zonation, gut lumen path) and transit times.

### Statistical Analysis

Graphpad Prism was used to conduct all statistical tests and 4-point logistic curve fitting for interpolation of fluorescent tracer concentrations from standard curves. Unless otherwise noted, p-values < 0.05 were considered significant, and mean and standard error of the mean (SEM) are shown in all plots.

## AUTHOR CONTRIBUTIONS

R.N., A.H., B.M.M., and R.P-B. led the data analysis for generation of figures and prepared the manuscript with D.E.I.; A.H., B.M.M., M.C., T.H., M.B., S.D., E.A.F., S.S.F.J., T.G., V.K., L.L., R.M., Y.M., J.N., B.O., T-E.P., H.S., B.S., G.J.T., Z.T., T.H-I., K-J.J., A.S.P and M.Y. planned and performed biological experiments under the supervision of G.A.H., O.L., A.B, R.N., R. P-B., K.K.P., and D.E.I.; A.C., E.C., Y.C., J.F., R.F., C.N., R.N., G.H., J.G., N.W, K.K.P., and D.E.I. were responsible for chip development and fabrication; J.S., G.T.II, C.H., J.F-A., J.A.G, and D.L conceptualized the robotics-based Organ Chip linking and developed early versions of the Interrogator instrument and software with an integrated robotic sampler; M.I., Y.C., S.C., A.D., T.D., T.F., O.H. and R.N were responsible for software and hardware engineering involved in the development of the final instrument and sensors used in this study; D.D., M.R.S., and A.P. were responsible for MCRO model development and data analysis working closely with A.H., B.M.M, R.P-B, and R.N.; and R.N., R.P-B., O.L., G.A.H., D.L., K.K.P. and D.E.I. were responsible for overseeing and orchestrating the entire effort.

## ACKNOWLEDGMENTS

This research was sponsored by the Wyss Institute for Biologically Inspired Engineering at Harvard University and the Defense Advanced Research Projects Agency under Cooperative Agreement Number W911NF-12-2-0036. The views and conclusions contained in this document are those of the authors and should not be interpreted as representing the official policies, either expressed or implied, of the Defense Advanced Research Projects Agency, or the U.S. Government. This work was performed in part at the Center for Nanoscale Systems (CNS), a member of the National Nanotechnology Coordinated Infrastructure Network (NNCI), which is supported by the National Science Foundation under NSF award no. 1541959. CNS is part of Harvard University, the Harvard Materials Research Science and Engineering Center (DMR-1420570). We thank John Caramanica and Paul Machado for their machining expertise, M. Rosnach for his artwork, B. Fountaine and S. Kroll for their help with photography, M. Rousseau for help with videography, C. Vidoudez for mass spectrometry analysis, and J. Wikswo for helpful input at the start of this project.

