## Supplementary Information for "A robotic platform for fluidically-linked human body-on-chips experimentation"

[1]Wyss Institute for Biologically Inspired Engineering at Harvard University, Boston, MA 02115, USA; [2]CFD Research Corporation, Huntsville, AL, USA 35806; [3]Harvard John A. Paulson School of Engineering and Applied Sciences, Harvard University, Cambridge, MA 02138, USA; [4]Department of Pathology, Massachusetts General Hospital, Boston, MA 02115, USA; [5]Department of Bioengineering and iBB - Institute for Bioengineering and Biosciences, Instituto Superior Técnico, Universidade de Lisboa, Lisboa, Portugal; [6]Vascular Biology Program and Department of Surgery, Boston Children's Hospital and Harvard Medical School, Boston, MA 02115, USA; [7]Department of Biology, University of Freiburg, Freiburg, Germany; [8]GlaxoSmithKline, 1250 S. Collegeville Rd., Collegeville, PA 19426, USA;

[*] Authors contributed equally

[+] Current address: Emulate, Inc., 27 Drydock Avenue, Boston, MA 02210, USA

**Stage Calibration**

The deck layout exists in three spaces: the physical layout, the CAD assembly and the web-based GUI. The calibration protocol developed here relates the positional error between the physical and CAD layouts. The measured positional errors can then be used to calculate linear scaling and offset values that, when applied in the web-based GUI, significantly reduce alignment error between the liquid handler and the individual deck components. A given x,y,z coordinate as described by the CAD layout is calibrated to the physical layout by multiplying the spatial coordinates by a calibration matrix and adding an offset matrix. A schematic of this calibration is shown in **Fig. 3b**. In a perfectly aligned system, the calibration matrix would simply be an identity matrix (Equation 1) and all offset values would be zero (Equation 2). In order to use this calibration procedure, robot alignment error must be determined by installing four precision ground post subassemblies (Thorlabs P series 1.5" diameter post, MiSUMi M6 10mm diameter locating pin) used as calibration references and a ruby-probe tip (Renishaw 1mm ball end 10 mm length, M4 thread) attached to the end of the liquid handler with a custom tapped 20 mm long spacer (M4 female to ¼-28 female). Two matrices are generated; matrix ‘P’, the theoretical, CAD-based, distance between the four measurement references (equation 3), and matrix ‘Q’, the physically measured distance between the four calibration references (equation 4). The web-based GUI features an interactive aid to complete the measurement process; upon completion, the calibration posts are removed. The process of determining the calibration and offset matrix is shown in equations 5 and 6, respectively. Once the M and O matrices have been determined, they are permanently stored within the client code and are applied to all motion system coordinates as directed by the web-based GUI. The calibration procedure is run when the system is assembled and re-calibrated upon software re-install, if any major changes are made to the deck layout, or at regularly scheduled maintenance intervals.

$$M = \begin{bmatrix} 1 & 0 & 0 \\ 0 & 1 & 0 \\ 0 & 0 & 1 \end{bmatrix} \quad \text{[Eq. 1]}$$

$$O = \begin{bmatrix} 0 & 0 & 0 \end{bmatrix} \quad \text{[Eq. 2]}$$

$$P = \begin{bmatrix} (B_{x1} - A_{x1}) & (B_{y1} - A_{y1}) & (B_{z1} - A_{z1}) \\ (C_{x1} - A_{x1}) & (C_{y1} - A_{y1}) & (C_{z1} - A_{z1}) \\ (D_{x1} - A_{x1}) & (D_{y1} - A_{y1}) & (D_{z1} - A_{z1}) \end{bmatrix} \quad \text{[Eq. 3]}$$

$$Q = \begin{bmatrix} (B_{x2} - A_{x2}) & (B_{y2} - A_{y2}) & (B_{z2} - A_{z2}) \\ (C_{x2} - A_{x2}) & (C_{y2} - A_{y2}) & (C_{z2} - A_{z2}) \\ (D_{x2} - A_{x2}) & (D_{y2} - A_{y2}) & (D_{z2} - A_{z2}) \end{bmatrix} \quad \text{[Eq. 4]}$$

$$M = [Q][P]^{-1} \quad \text{[Eq. 5]}$$

$$[O_x \quad O_y \quad O_z] = [A_{x2} \quad A_{y2} \quad A_{z2}] - [A_{x1} \quad A_{y1} \quad A_{z1}]\,[M] \quad \text{[Eq. 6]}$$

**SI Table 1. Organ Chip types and associated properties of the microfluidic features.**

| Organ Chip | Membrane | Channel Width (mm) | Apical Channel Height (mm) | Basal Channel Height (mm) | Channel Length (mm) |
|---|---|---|---|---|---|
| Gut | PDMS 7 µm pores | 1 | 1 | 0.2 | 91 |
| Liver | PDMS 7 µm pores | 1 | 1 | 0.2 | 24 |
| Kidney | PET 0.4 µm pores | 1 | 0.1 | 0.1 | 24 |
| Heart | PC 5 µm pores | 2.5 | 0.25 | 1.2 | 25 |
| Lung | PDMS 7µm pores | 1 | 1 | 0.2 | 24 |
| Skin | PDMS 7µm pores | 5.5 (apical)<br>1 (basal) | 5 | 0.3 | 24 |
| BBB | PET 0.4 µm pores | 1 | 1 | 0.2 | 24 |
| Brain | PC 5 µm pores | 2.5 | 0.1 | 1.2 | 25 |

PDMS: poly (dimethyl siloxane); PET: polyester terephthalate; PC: polycarbonate

**SI Table 2 – Cell culture parameters**

| Organ | Chip Type | ECM Coating | Apical | | Basal | |
|---|---|---|---|---|---|---|
| | | | **Cell Type** | **Seeding Density** | **Cell Type** | **Seeding Density** |
| **Gut** | Long Tall Channel, Stretchable, PDMS Membrane | Matrigel & Collagen I | Caco2 BBE | 100,000 cells/cm$^2$ | HUVEC | 100,000 cells/cm$^2$ |
| **Liver** | Standard Tall Channel, Stretchable, PDMS Membrane | Collagen I | Primary Human Hepatocytes | 250,000 cells/cm$^2$ | LSEC | 100,000 cells/cm$^2$ |
| **Kidney** | Standard tall channel, PET membrane | Collagen IV & Laminin | Human Renal Proximal Tubule | 120,000 cells/cm$^2$ | Human Glomerular Microvascular Endothelial | 100,000 cells/cm$^2$ |
| **Heart** | Dual channel PC, PET Membrane | Fibronectin | Human iPSC Cardiomyocytes (Cor4U, Axiogenesis) | 200,000 cell/cm$^2$ | HUVEC | 100,000 cells/cm$^2$ |
| **Lung** | Standard Tall Channel, Stretchable, PDMS Membrane | Fibronectin & Collagen I | A549 Human Lung Carcinoma Cell Line | 200,000 cells/cm$^2$ | HUVEC | 60,000 cells/cm$^2$ |
| **BBB** | Standard tall channel, PET membrane | Collagen IV & Fibronectin | Primary Human Astrocytes | 70,000 cells/cm$^2$ | Primary Human Brain Microvascular Endothelial | 90,000 cells/cm$^2$ |
| | | | Primary Human Pericytes | 30,000 cells/cm$^2$ | | |
| **Brain** | Dual channel PC, PET Membrane | PDL & Laminin | Human Hippocampal Neural Stem Cells (hNSCs) | 100,000 cells/cm$^2$ | N/A | |
| **Skin** | Oval Open-top, Stretchable, PDMS Membrane | Fibronectin & Collagen I | Human Primary Adult Dermal Fibroblasts | 82,000 cells/cm$^2$ | Human Primary Dermal Microvascular Endothelial | 100,000 cells/cm$^2$ |
| | | | Human Primary Neonatal Epidermal Keratinocytes | 800,000 cells/cm$^2$ | | |

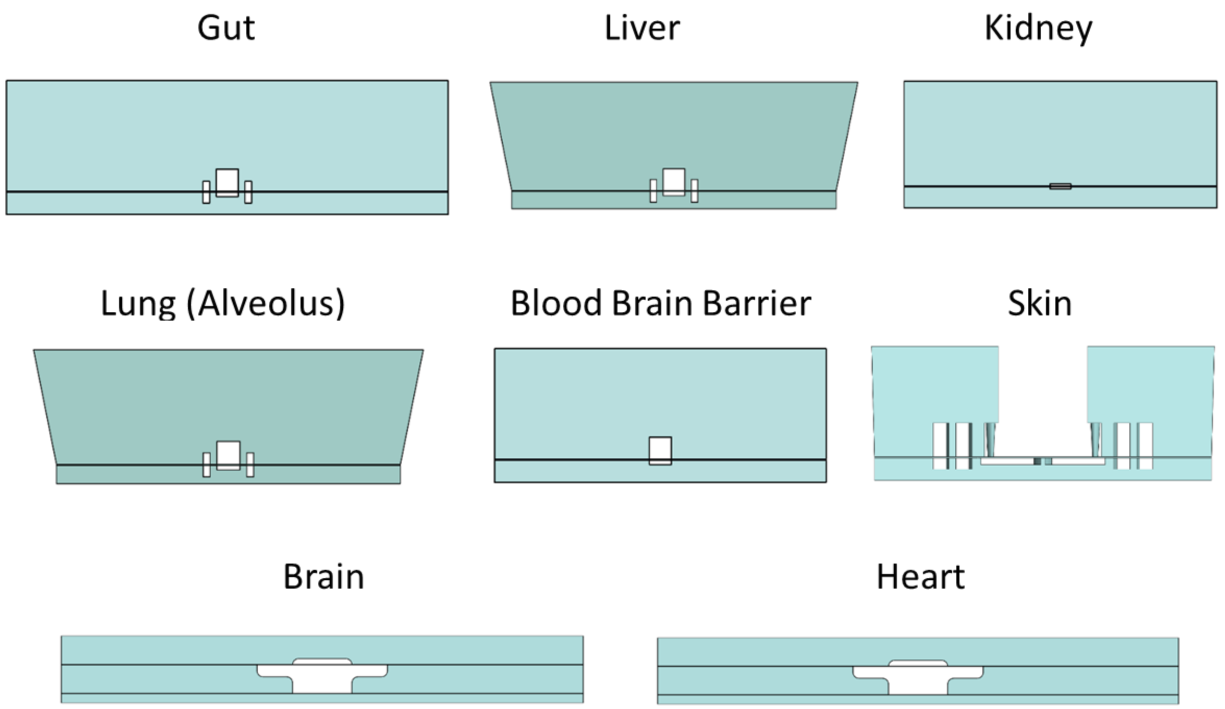


**Supplementary Figure S1.** Cross sectional views of Organ Chip CAD for all 8 Organ Chips used in this study. In all chips, two parallel fluidic channels are separated by a porous semi-permeable membrane.

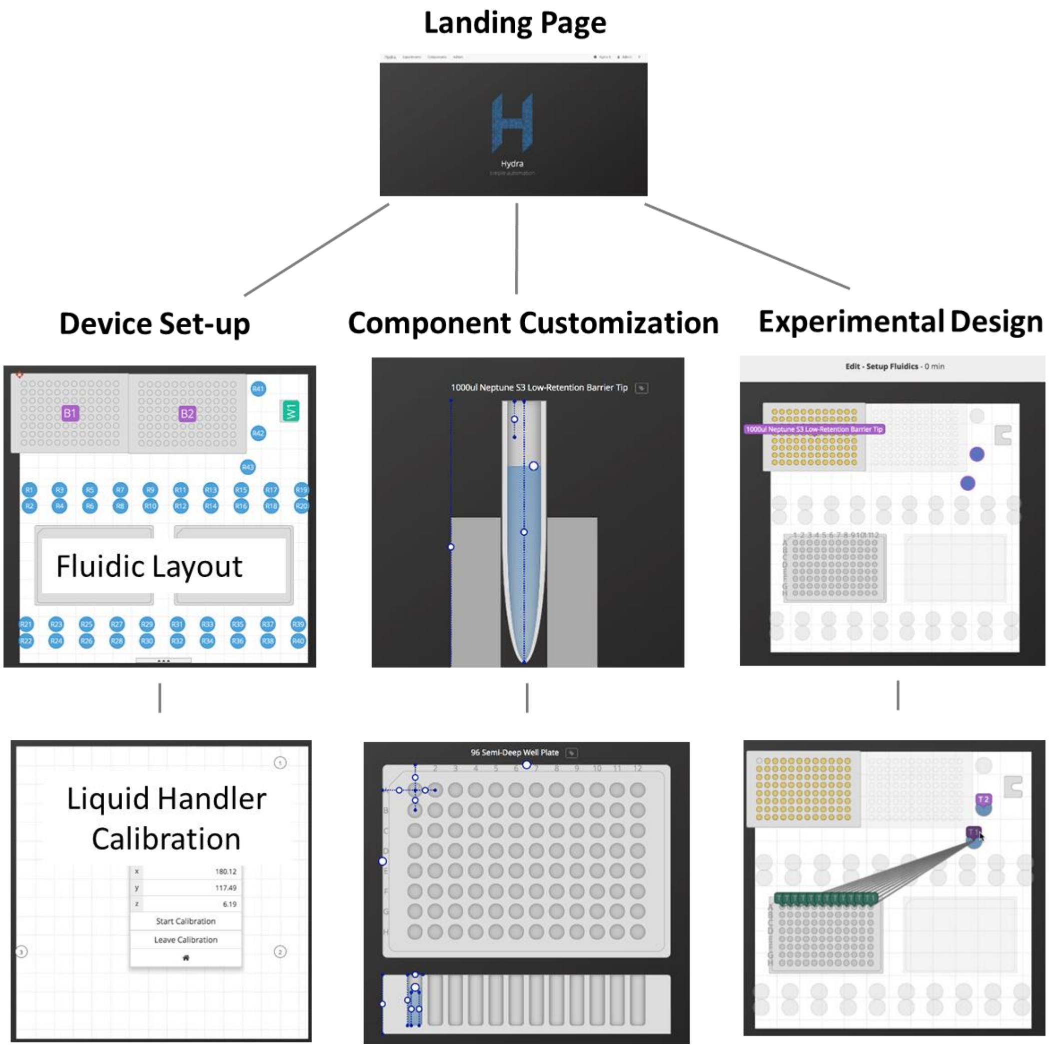


**Supplementary Figure S2.** Hydra software screen shots highlighting key features of device setup, component customization, and experimental design. All Interrogator programming is performed using this graphical Hydra software interface.

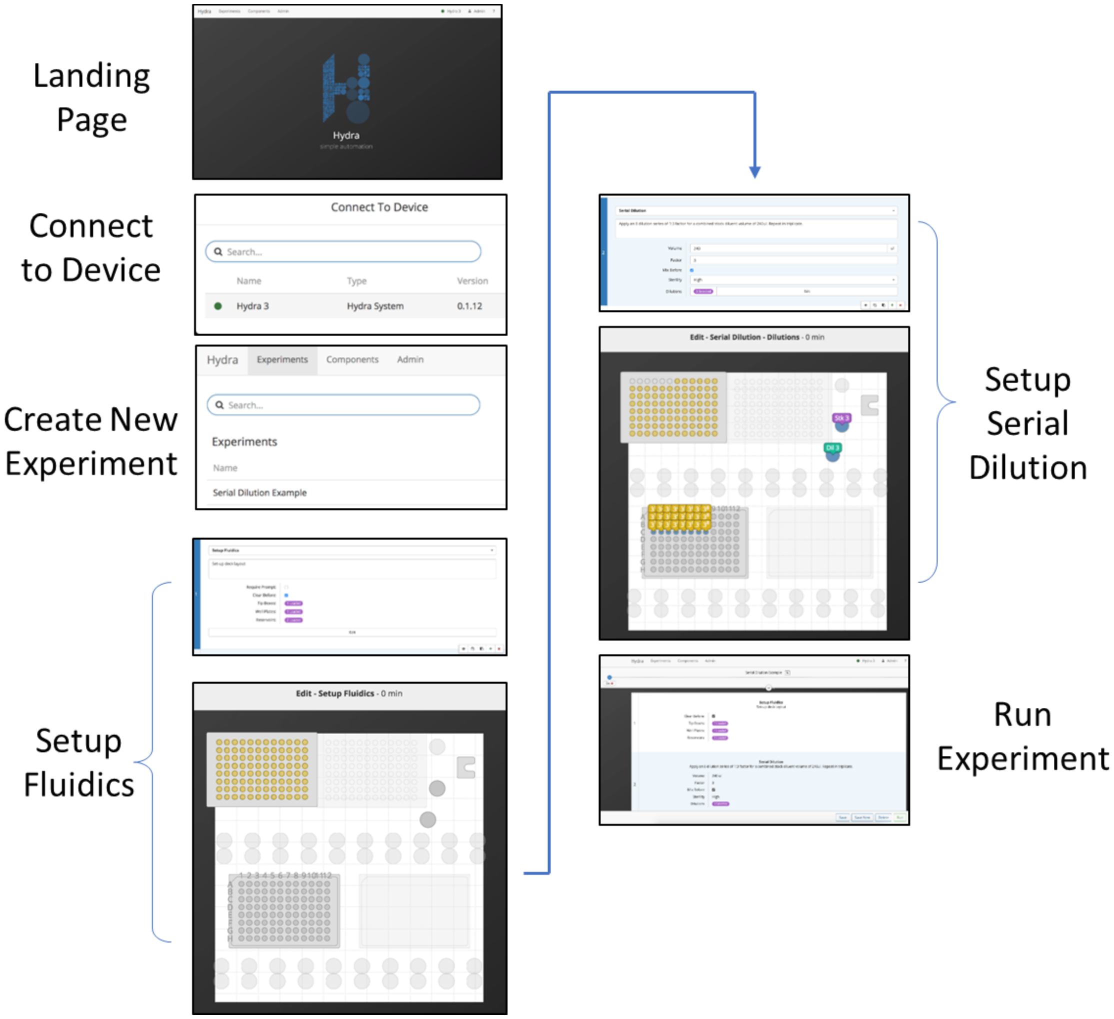


**Supplementary Figure S3.** Hydra software screen shots of the rapid workflow involved in designing and executing a typical experiment

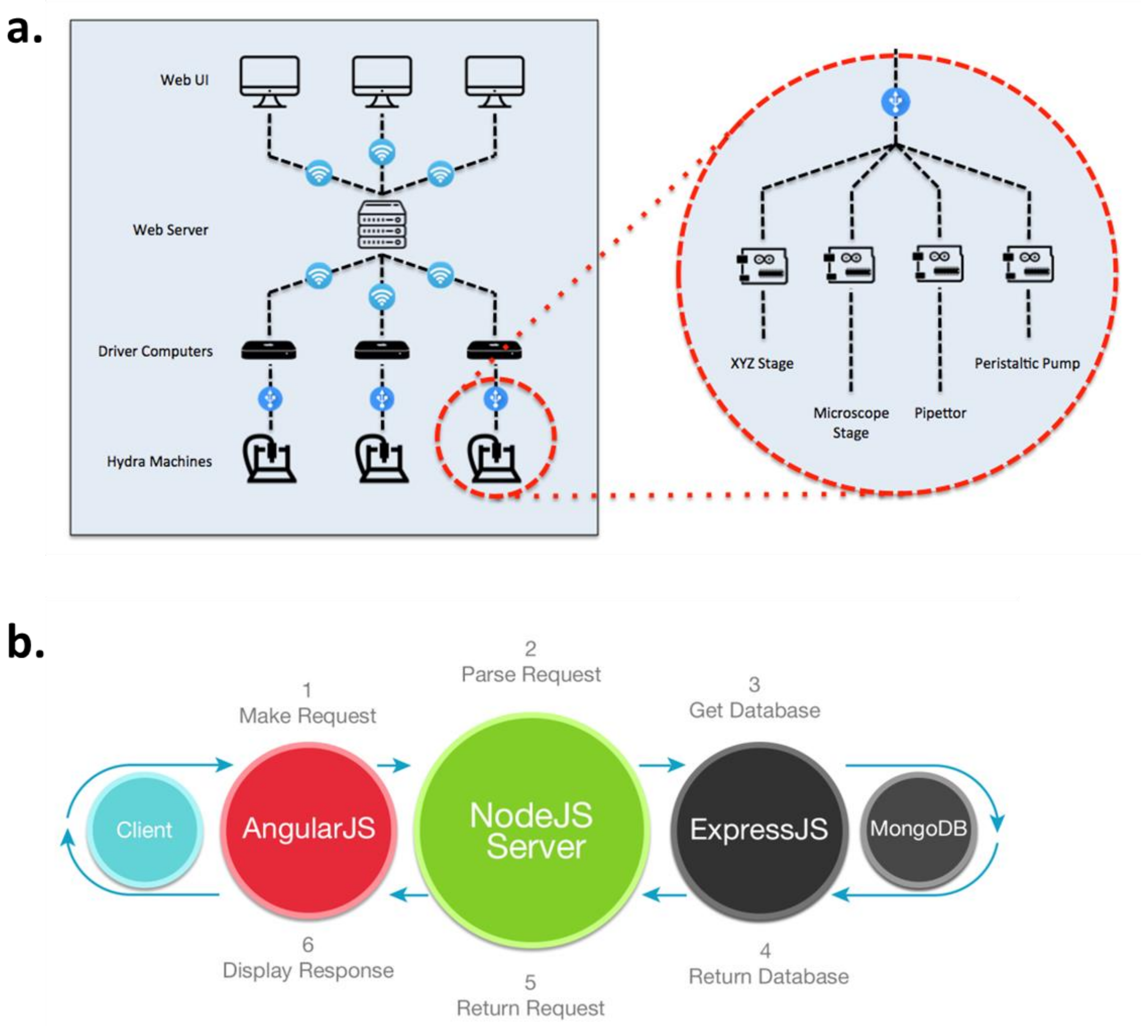


**Supplementary Figure S4.** Schematic of Hydra (A) system communication structure and (B) network software architecture.

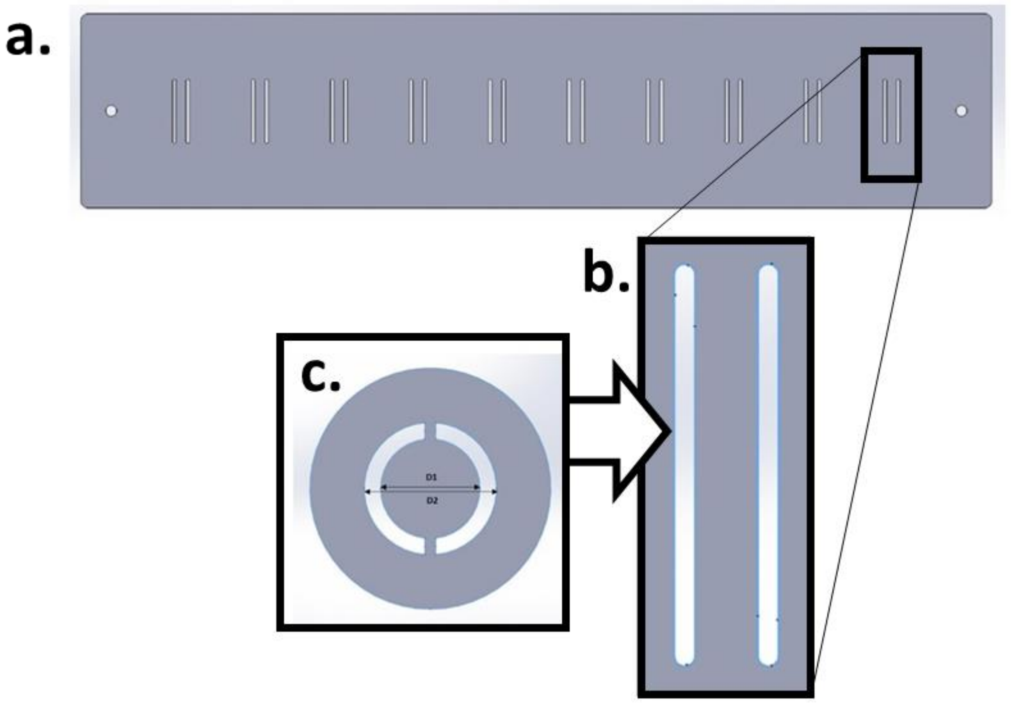


**Supplementary Figure S5.** Rendering of the phase plate (A) used to generate phase contrast-like illumination in Organ Chips while enabling scanning across their channel length. Each Organ Chip cartridge location is directly below each phase slot feature (B), which was designed based on standard condenser phase ring geometry (C).

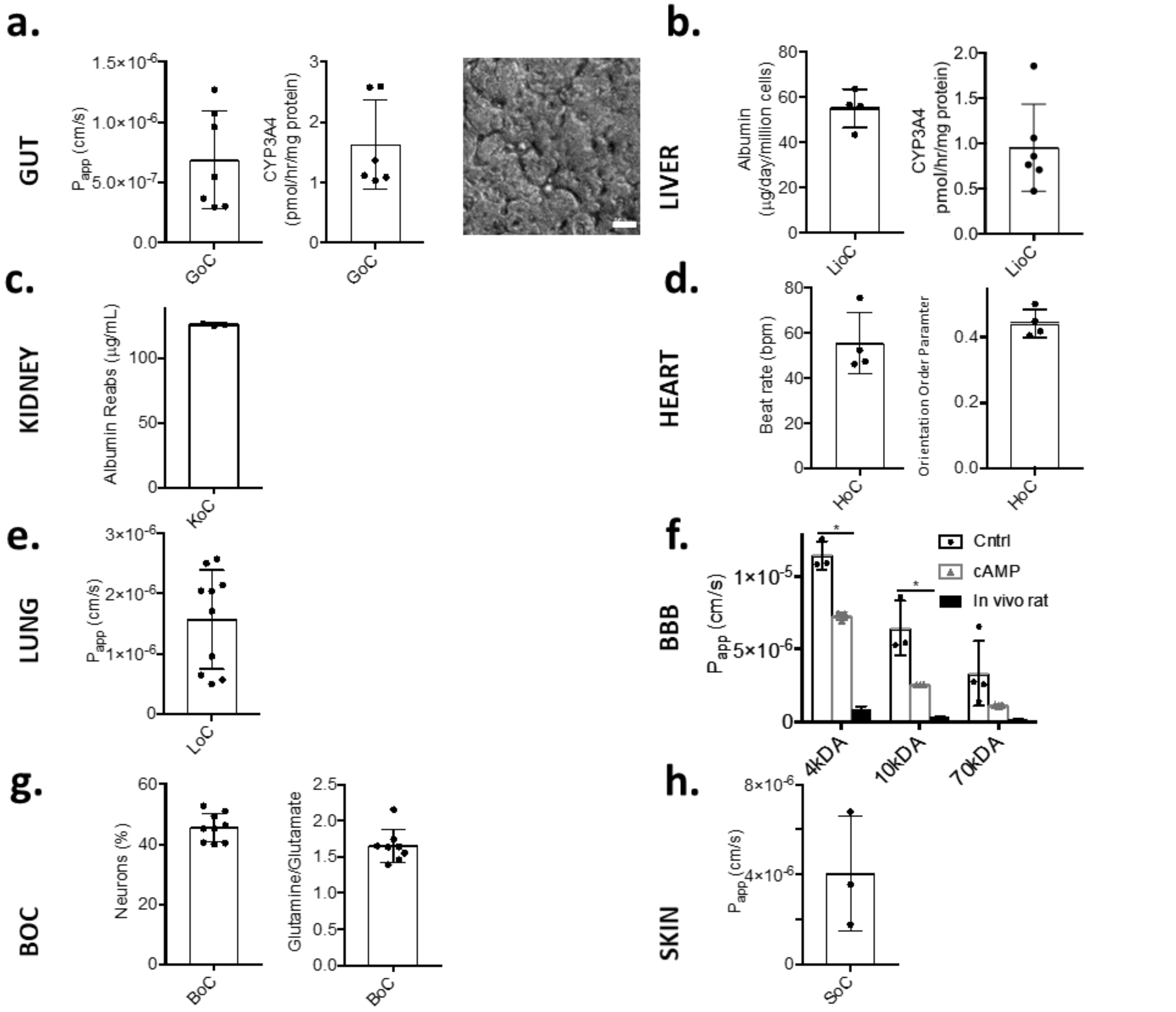


**Supplementary Figure S6.** Organ Chip-specific readouts prior to linking demonstrating organ maturity. (A) Gut Chip: Cascade blue® (596 Da) permeability, n=7, CYP3A4 activity, and bright field imaging of villi, scale 50 μm. (B) Liver Chip: albumin production and CYP3A4 activity, n=4 and n=6, influence of liver endothelial cells on Liver Chip function, n=2 (C) Kidney Chip: albumin reabsorption, n=3 (D) Heart Chip: beat rate and orientation order parameter (OOP[11]), n=4 (E) Lung Chip: Texas Red™ (3 kDa) permeability, n=10 (F) BBB Chip permeability effect of Cyclic adenosine monophosphate (cAMP) and comparison to rat *in vivo* data : Inulin-FITC (2-5 kDa, listed as 4 kDa for comparison with literature *in vivo* data) p=0.002, n=3, Dextran-Cascade blue® (10 kDa) p= 0.024, n=3, Dextran-Texas Red™ (70 kDa) permeability, n=4, *in vivo* data[12] (G) Brain Chip: ratio of neuronal cells and glutamine:glutamate ratio, n=9 and n=8 (H) Skin Chip: Cascade Blue (596Da) permeability, n=3.

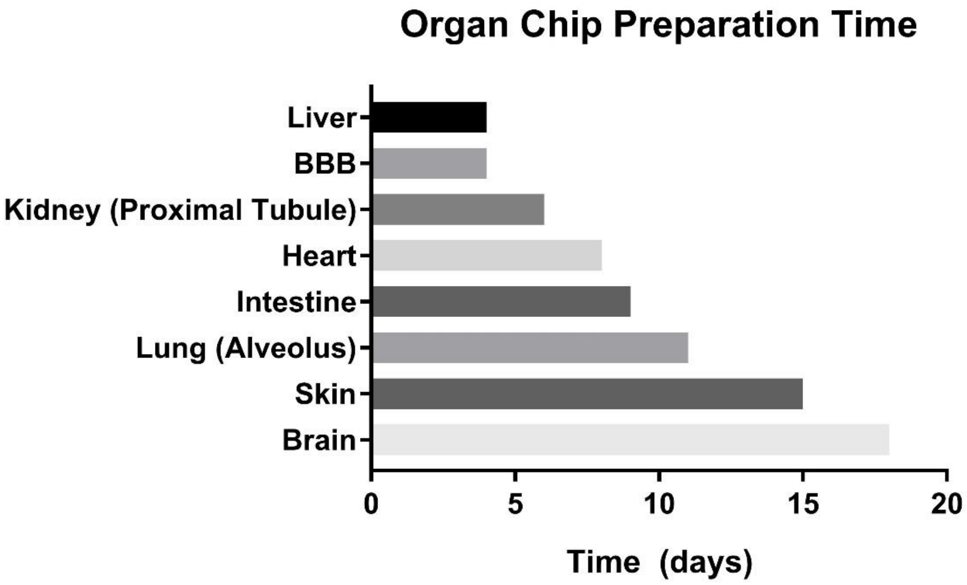


**Supplementary Figure S7.** Length of culture time required for Organ Chips to reach a mature morphological and functional state. After the indicated time the Organ Chips were used for linkage experiments.

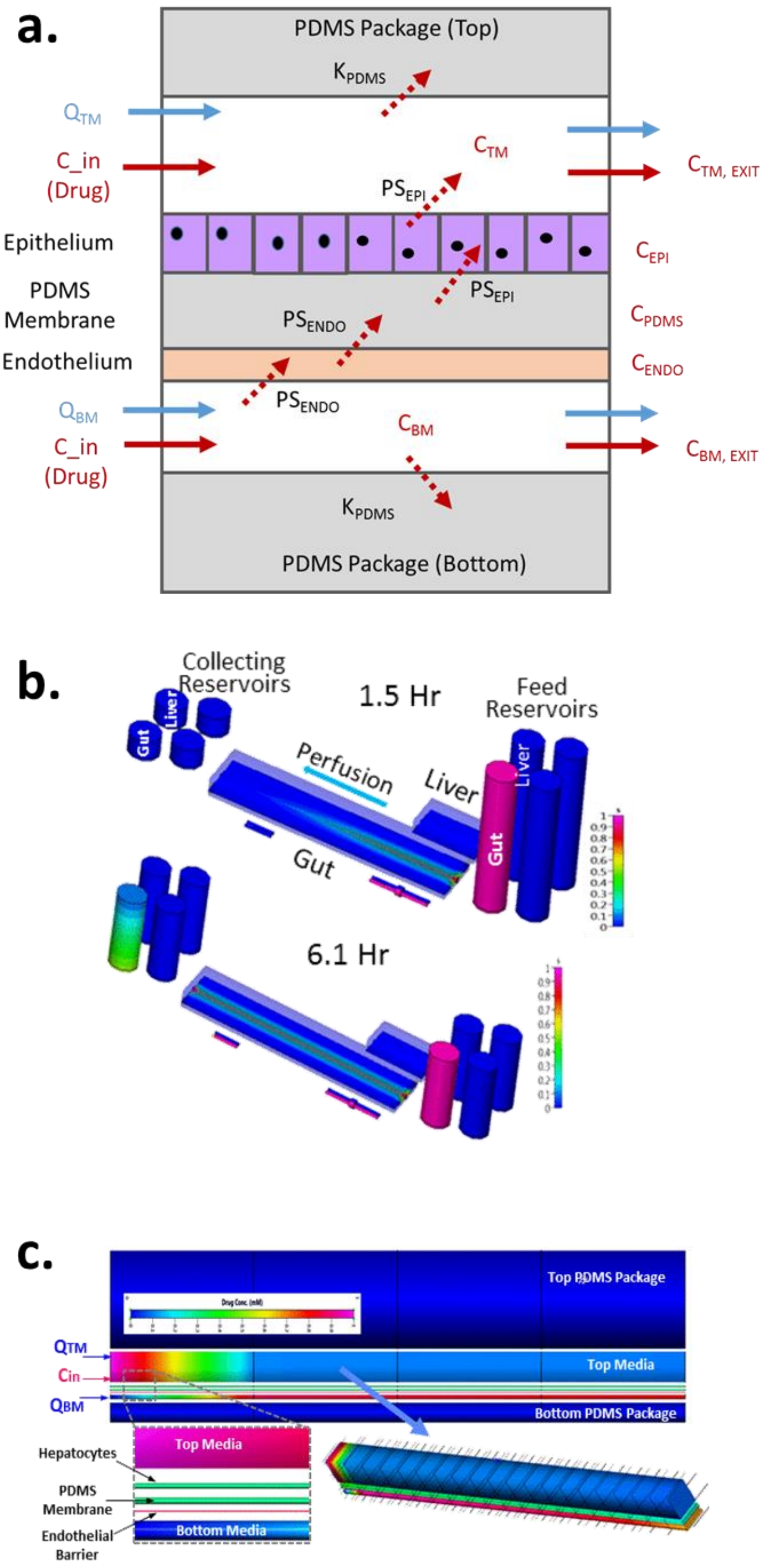
a.
PDMS Package (Top)
$K_{PDMS}$
$Q_{TM}$
C_in
(Drug)
$C_{TM}$
$C_{TM, EXIT}$
$PS_{EPI}$
Epithelium
$C_{EPI}$
PDMS
Membrane
$PS_{EPI}$
$C_{PDMS}$
$PS_{ENDO}$
Endothelium
$C_{ENDO}$
$Q_{BM}$
$PS_{ENDO}$
$C_{BM}$
C_in
(Drug)
$C_{BM, EXIT}$
$K_{PDMS}$
PDMS Package (Bottom)
b.
Collecting
Reservoirs
Gut
Liver
1.5 Hr
Feed
Reservoirs
Perfusion
Liver
Liver
Gut
Gut
6.1 Hr
c.
Top PDMS Package
$Q_{TM}$
Cin
$Q_{BM}$
Top Media
Bottom PDMS Package
Hepatocytes
Top Media
PDMS
Membrane
Endothelial
Barrier
Bottom Media

**Supplementary Figure S8.** Computational Microphysiological Model of a generic Organ Chip (A). Each Organ Chip model is spatially resolved in both stream-wise and cross-stream direction (top PDMS, top channel media, epithelial cell layer, membrane, endothelial cell layer, bottom channel media, bottom PDMS). (B) A quantitative reduced order 3D model (Q3D) enables tracking drug dynamics through linked Organ Chips (Gut to Liver shown here). Two time instances show initial transport of the gut microchannel at 1.5 hr and at 6.1hr. The media from “vascular” collecting reservoirs is periodically transferred to downstream linked Organ Chips using the Interrogator. (C) Detailed view of the 3D spatially resolved Organ Chip model showing the geometrical/mesh resolution in the cross-stream and stream-wise (axial) directions.

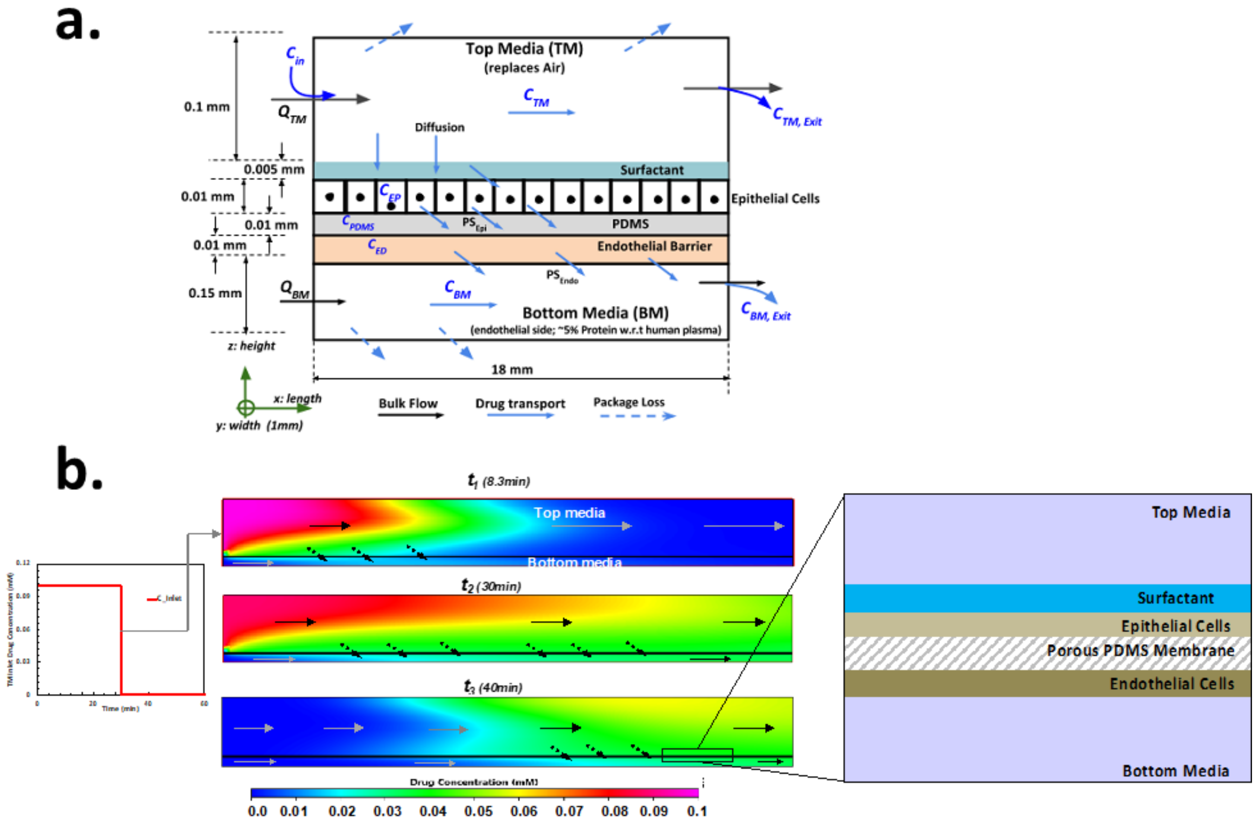


**Supplementary Figure S9.** Computational Model of the Lung Chip (A) Reduced-Order Model of Lung Chip in liquid-liquid interface culture, 4 compartments in axial direction, (B) High-Fidelity Model of drug bolus movement and apical→basal mass transfer through Lung Chip membrane over time.

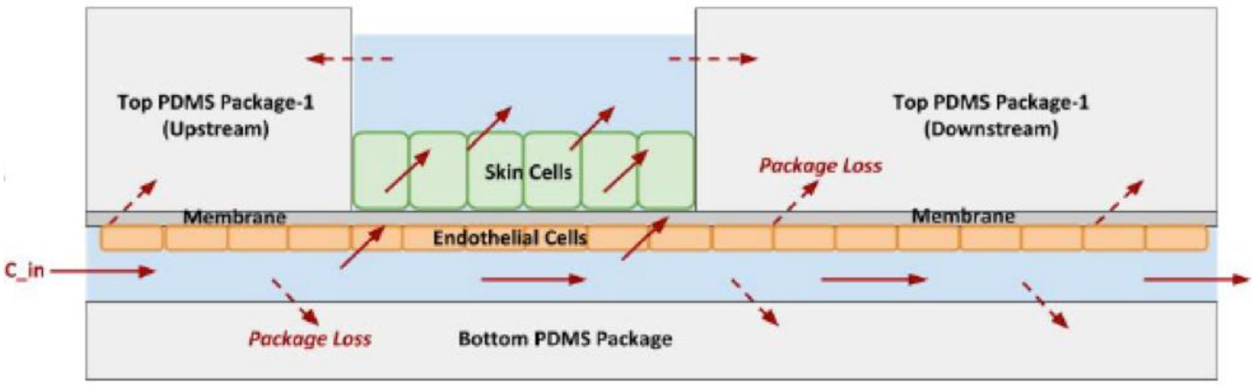


**Supplementary Figure S10.** Computational Model of the Skin Chip: Reduced-Order Model of Skin Chip, 4 compartments in axial direction

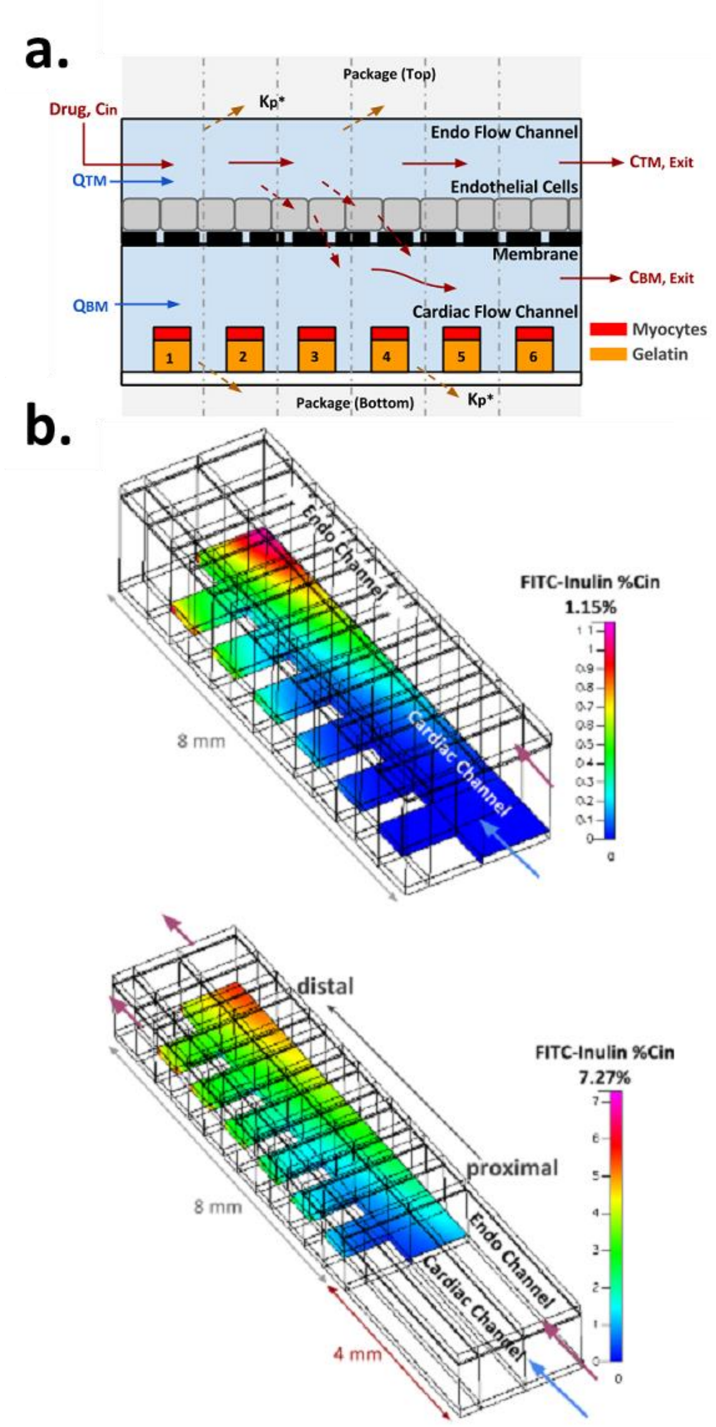


**Supplementary Figure S11.** Computational model of the Heart Chip A) Reduced-Order Model of MTF Heart Chip (6 compartments in axial direction), (B) High-fidelity models of MTF Heart Chip predict increased apical→basolateral mass transfer of high MW tracer when increasing chip length from 8 mm (top) to 12 mm (bottom).

a.

| Parameter | Value | Unit |
|---|---|---|
| Step Pulse | 10 | µs |
| Step Idle Delay | 100 | ms |
| Step Size (X & Y, Z) | 20.051, 200.000 | step/mm |
| Max Velocity (X & Y, Z) | 200, 56 | mm/s |
| Acceleration (X & Y, Z) | 400, 160 | mm/s$^2$ |
| Motion Envelope (X & Y, Z) | 310, 135 | mm |

b.

```
$0=10 (step pulse, usec)
$1=100 (step idle delay, msec)
$2=0 (step port invert mask:00000000)
$3=3 (dir port invert mask:00000011)
$4=0 (step enable invert, bool)
$5=1 (limit pins invert, bool)
$6=0 (probe pin invert, bool)
$10=1 (status report mask:00000001)
$11=0.020 (junction deviation, mm)
$12=0.010 (arc tolerance, mm)
$13=0 (report inches, bool)
$20=1 (soft limits, bool)
$21=1 (hard limits, bool)
$22=1 (homing cycle, bool)
$23=7 (homing dir invert mask:00000111)
$24=60.000 (homing feed, mm/min)
$25=2000.000 (homing seek, mm/min)
$26=100 (homing debounce, msec)
$27=1.000 (homing pull-off, mm)
$100=20.051 (x, step/mm)
$101=20.051 (y, step/mm)
$102=200.000 (z, step/mm)
$110=12000.000 (x max rate, mm/min)
$111=12000.000 (y max rate, mm/min)
$112=3400.000 (z max rate, mm/min)
$120=400.000 (x accel, mm/sec^2)
$121=400.000 (y accel, mm/sec^2)
$122=160.000 (z accel, mm/sec^2)
$130=310.000 (x max travel, mm)
$131=310.000 (y max travel, mm)
$132=135.000 (z max travel, mm)
```

**Supplementary Figure S12.** Motion system parameters (A) and GRBL shield settings (B) that define the characteristic behavior of the system.

### Supplementary Movies

**SI Movie 1**

**Overview of Interrogator system components and Organ Chip linking**.

**SI Movie 2**

**Microscope module movie of gut chip stretching.**

**SI Movie 3**

**Heart Chip beating after being linked for 3 weeks on the Interrogator platform.**

**SI Movie 4**

**High-fidelity diffusion model of the Heart Chip**.

**SI Movie 5**

**CoBi Q-3D Model of Gut-Liver Chip Linking**
