## Supplementary figures and images for "A robotic platform for fluidically-linked human body-on-chips experimentation"

### SI Movie 5

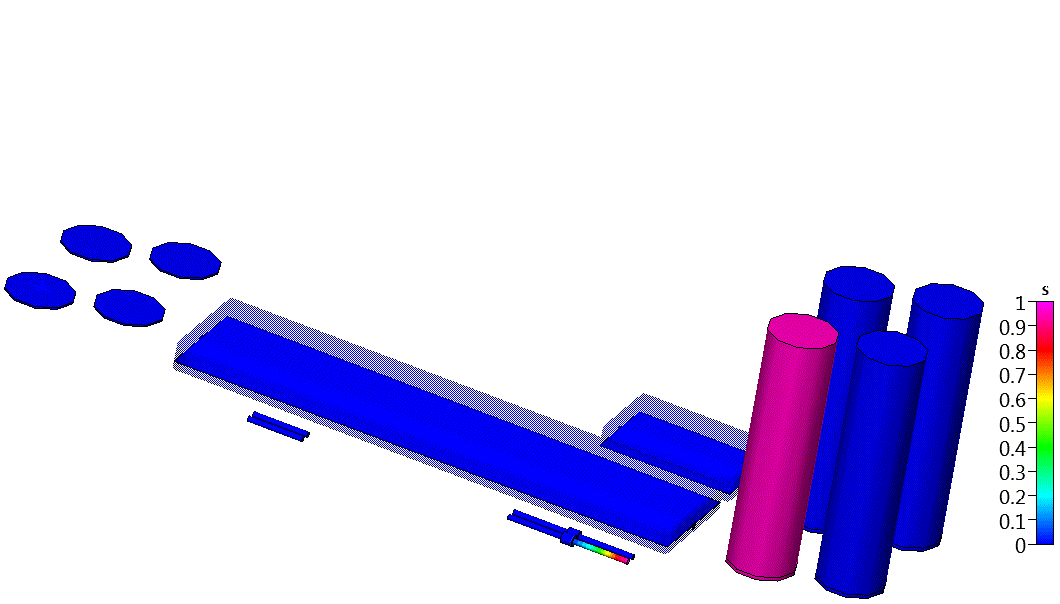
